# Evaluation of Vicinity-based Hidden Markov Models for Genotype Imputation

**DOI:** 10.1101/2021.09.28.462261

**Authors:** Su Wang, Miran Kim, Xiaoqian Jiang, Arif Harmanci

## Abstract

The decreasing cost of DNA sequencing has led to a great increase in our knowledge about genetic variation. While population-scale projects bring important insight into genotype-phenotype relationships, the cost of performing whole-genome sequencing on large samples is still prohibitive. In-silico genotype imputation coupled with genotyping-by-arrays is a cost-effective and accurate alternative for genotyping of common and uncommon variants. Imputation methods compare the genotypes of the typed variants with the large population-specific reference panels and estimate the genotypes of untyped variants by making use of the linkage disequilibrium patterns. Most accurate imputation methods are based on the Li-Stephens hidden Markov model, HMM, that treats the sequence of each chromosome as a mosaic of the haplotypes from the reference panel. Here we assess the accuracy of local-HMMs, where each untyped variant is imputed using the typed variants in a small window around itself (as small as 1 centimorgan). Locality-based imputation is used recently by machine learning-based genotype imputation approaches. We assess how the parameters of the local-HMMs impact the imputation accuracy in a comprehensive set of benchmarks and show that local-HMMs can accurately impute common and uncommon variants and can be relaxed to impute rare variants as well. The source code for the local HMM implementations is publicly available at https://github.com/harmancilab/LoHaMMer.

## Introduction

As the cost of DNA sequencing is decreasing, the number of available genome sequences is increasing at a fast pace^1–4^. DNA sequencing is also the fundamental step for technologies such as RNA sequencing and ChIP-Sequencing^5^. Currently, there are millions of genomic sequences available and many more are expected^6–8^. As the genomic data is used more prevalently in the clinic and in translational research^9,10^, the genetic data size is available in many different scenarios, even including the citizen scientists from the general population^11^. Genetic data is deposited widespread in many places (including personal computers and even phones) and it made its way well into the fields of recreational genetics^12^. This is made possible by extensive mapping of the genetic differences between populations and efficient methods that can sift through massive databases for searching for relatives^13^. These are made possible by population-scale projects such as UKBiobank^14^. The third parties aim at capitalizing on these data by generating “new and interesting” datasets from rare phenotypes, diseases, and rare populations and individuals^15–18^. The genetic data is treated as a new type of commodity through brokers^19,20^. As the COVID19 pandemic showed, sharing and using genetic data out of the research lab and in the field and clinic will be routine tasks in many cases^21,22^.

One of the main uses of genetic data is performing genotype-phenotype associations using genome-wide association studies (GWAS or GWA study)^23–26^. For this, a large cohort is generated and the individuals are genotyped by sequencing. Next, the phenotype of interest (Intelligence quotient, height, body-mass index, blood glucose levels, etc.) is measured from all the individuals. Finally, the measured genotypes for all the variants are tested for association with the GWA studies, most variants are found to be in intergenic regions out of the protein-coding exons. Thus, it is necessary to perform genotyping on the whole genome scale using, for example, whole-genome sequencing (WGS) to ensure that the causal variants, not another variant that is in linkage disequilibrium (LD)^27,28^, can be identified as accurately as possible. This, however, is not cost-effective because large samples must be whole-genome sequenced^29^. To get around this, genotyping arrays are used for genotyping and decreasing the cost^30^. The genotyping arrays are designed to genotype only a sparse set of variants from the genome. These variants are then input to in-silico genotype imputation algorithms^31,32^, which impute and “fill-in” the un-genotyped (or untyped for short) variants. The main idea behind the imputation algorithms is to make use of the known haplotype structure of the whole genome and estimate the genotypes of the untyped variants using the genotypes of typed-variants that are correlated at the haplotype level^33^. The haplotype structure arises because the alleles are inherited between generations by a limited number of crossing-overs at the recombination hotspots between homologous chromosomes^34^. This causes long chunks of haplotypes to be inherited as a single unit. Although the length of conserved chunks (identity-by-descent segments^35^) decreases as the relationship distance increases, it can still be detected even with 20-25 generations of separation between individuals^36,37^. The imputation algorithms focus on making use of conserved chunks of haplotypes (i.e., frequent haplotypes) that are shared among unrelated individuals in the population. Imputation methods are also used for imputing variants identified by the RNA sequencing and whole-exome sequencing and for fine mapping of the variants from association studies.

The current state-of-the-art imputation methods such as BEAGLE, Minimac, and IMPUTE suite make use of the hidden Markov model (HMM)^38,39^ based approach that is developed by Li and Stephens^40–45^. HMM treats each haplotype as a “state” and analyzes the probabilities of all the “paths” that pass through the states to generate the alleles that are typed by the array^41^. This way, HMM-based methods can assign probabilities to the imputed genotypes using the probabilistic model imposed by the Li-Stephens haplotype model. The HMM takes the typed variants and the reference panel as input and imputes all the variants that exist on the reference panel but are untyped by the genotyping array. While HMM models provide good accuracy of imputation, they may fail at imputation of rare variants as these variants are represented on at least as rare haplotypes^46^. However, as the size of reference panels increases, the rare variants can be more accurately predicted^47^.

Here, we focus on Li-Stephens HMM-based imputation models and assess the performance of “local-HMMs”, i.e., the HMM evaluates the paths over only a short stretch of variants around the untyped variants. While several methods have tested different parametrizations of the state-of-the-art methods, we implemented the local-HMM methods to have full control over how the parameters impact the imputation. The locality-based approaches have been used in different scenarios, for instance with linear imputation models and with Deep Learning-based imputation models^48–50^, where the imputation is performed on the typed variants that are in the vicinity of the untyped variant. Also, IMPUTE and BEAGLE make use of a sliding-window, as long as 40 centimorgans (cM) to cut corners in computation. This parameter was not extensively assessed in terms of its impact on imputation accuracy. We implemented the per-position posterior probability estimation (we refer to this as the MAP method) by the forward-backward algorithm. We also implement the inference of the maximum-likelihood HMM path (referred to as the ML method), which represents the most likely mosaic of reference haplotypes that gives rise to the genotypes of the typed variants. On these methods, we analyze the size of the window, positioning of the target within the window, number of tag variants on the window, and the effective population size. It should be noted that we focus on the phased genotype imputation problem, i.e., we assume that the genotypes are phased. This is a reasonable assumption since pre-phasing has been showed to improve the time complexity of the imputation method substantially while conferring a very small performance penalty^44^.

One of the main advantages of the locality-based approaches is that they can be constrained in terms of computational requirements without the need to load the whole genome into memory or running the HMM inference methods on whole chromosomes. This way, the architecture of imputation algorithms on a cloud can be structured accordingly, for example, by using different models in different parts of the genome. On another front, the recent development of privacy-aware genotype imputation methods make use of the vicinity-based models and therefore our study can inform these methods about the locality parameters that must be considered and evaluated for increasing the imputation accuracy while providing privacy and confidentiality to the genetic data. The implementation of the local-HMMs, named LoHaMMer, is publicly available to download from https://github.com/harmancilab/LoHaMMer.

## Results

We briefly describe the HMM-based imputation techniques, the parameters, and the evaluation approach. We finally present the imputation accuracy evaluations.

### Overview of the Vicinity-based HMMs

Genotype imputation is summarized in Figure 1. The genotype imputation process takes as input the variant phased genotypes matrix, *G*_*M*×*V*_, individuals. As we are evaluating the phased imputation process, *G* is pre-phased using a phasing algorithm such as Eagle^51^. *G*_*i,j*_ holds the phased genotype of the *j*^*th*^ variant for the *i*^*th*^ individual, i.e 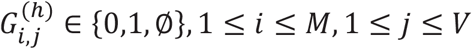, where *h* indicates the paternal/maternal copy for the genotype, i.e. 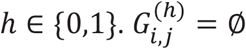 indicates the missing genotype that will be imputed using the reference panel. We denote the set of indices of the untyped variants with 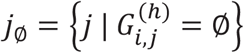. Imputation also takes the reference genotype matrix containing *H*_*N*×*V*_ of *N* haplotypes over the same *V* variants that correspond to the columns of *G*. Similar to *G, H*_*i,j*_ ∈ {0,1}, 1 ≤ *i* ≤ *N*, 1 ≤ *j* ≤ *V*.

**Figure 1.**
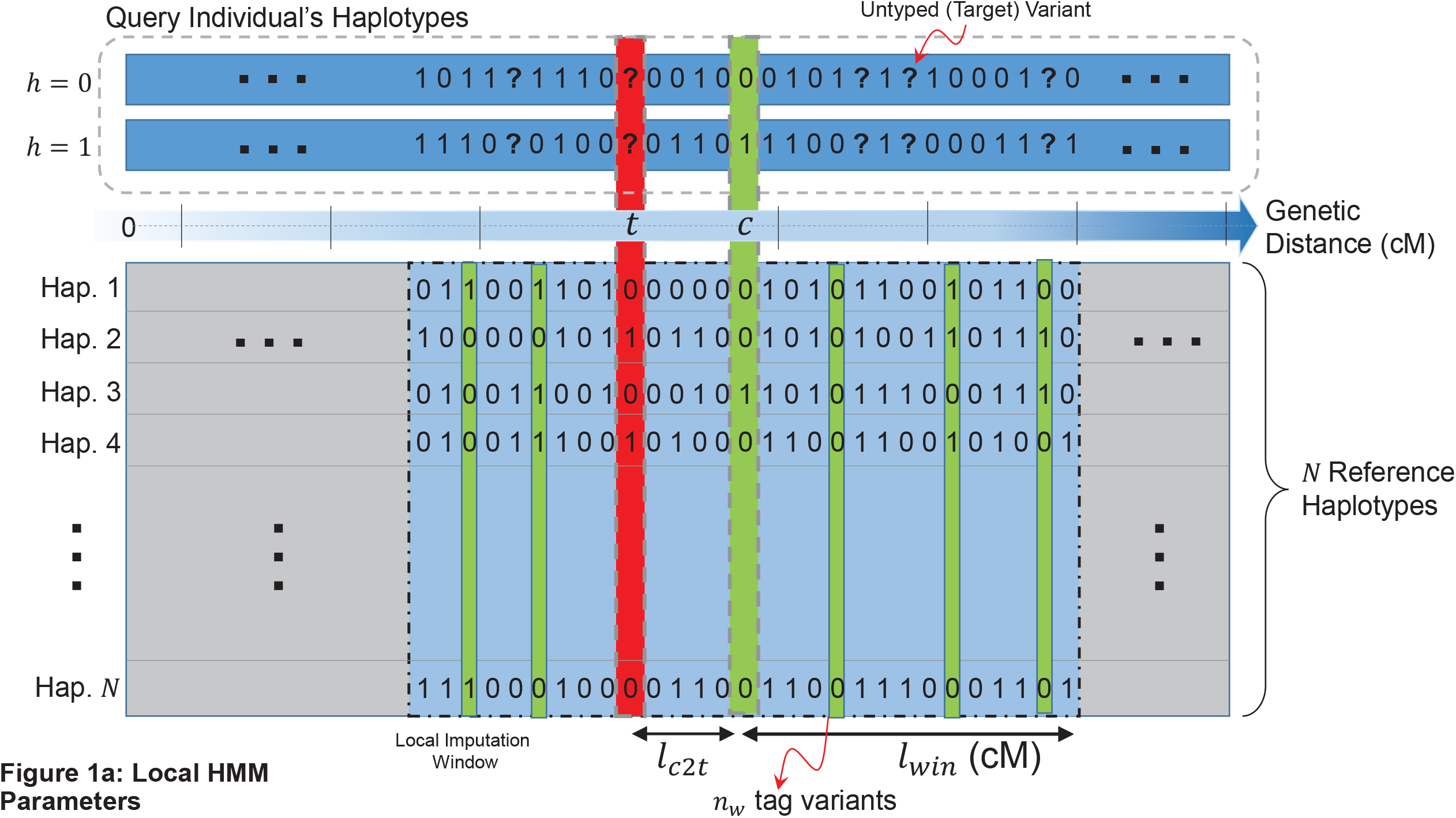
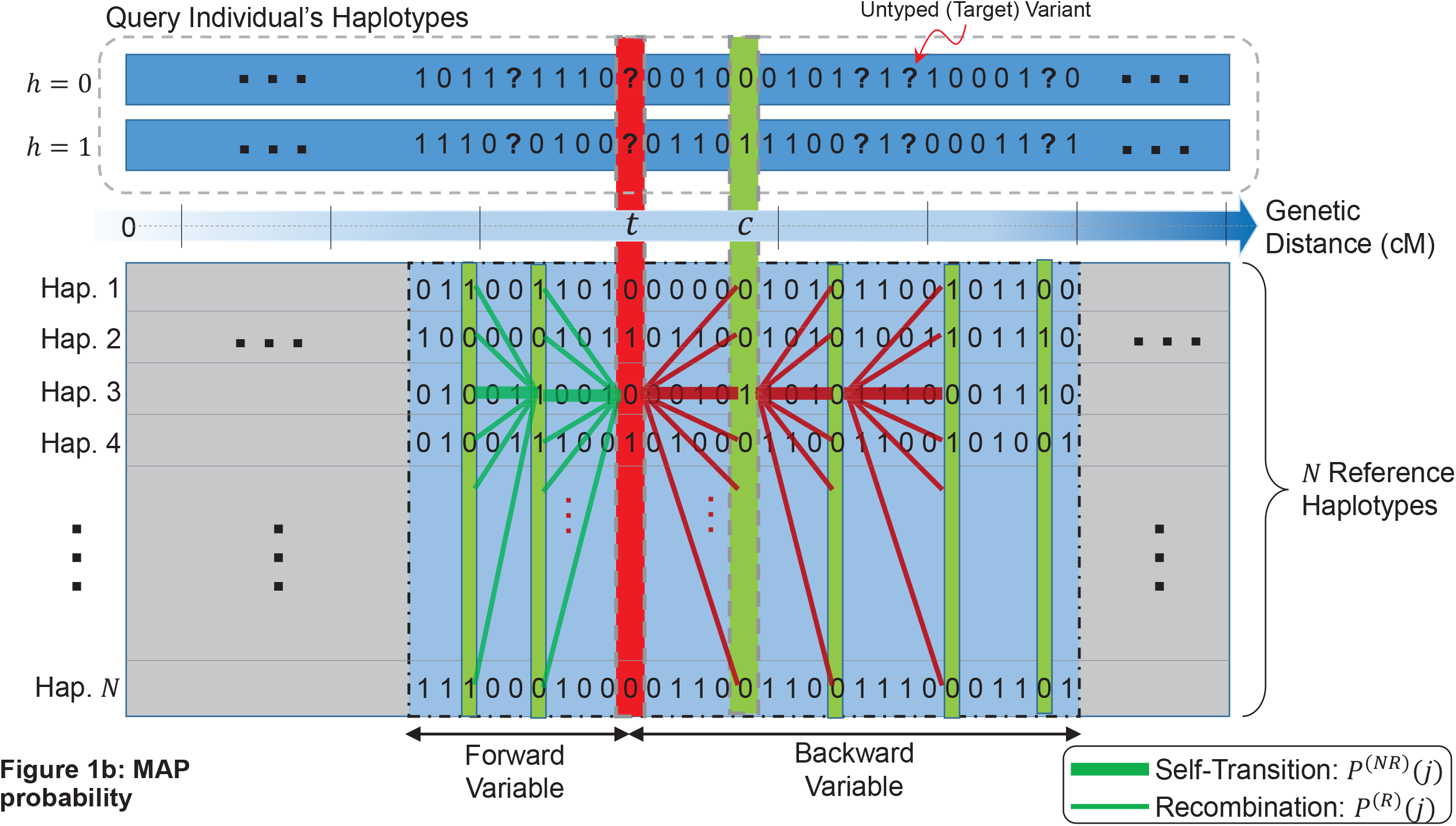
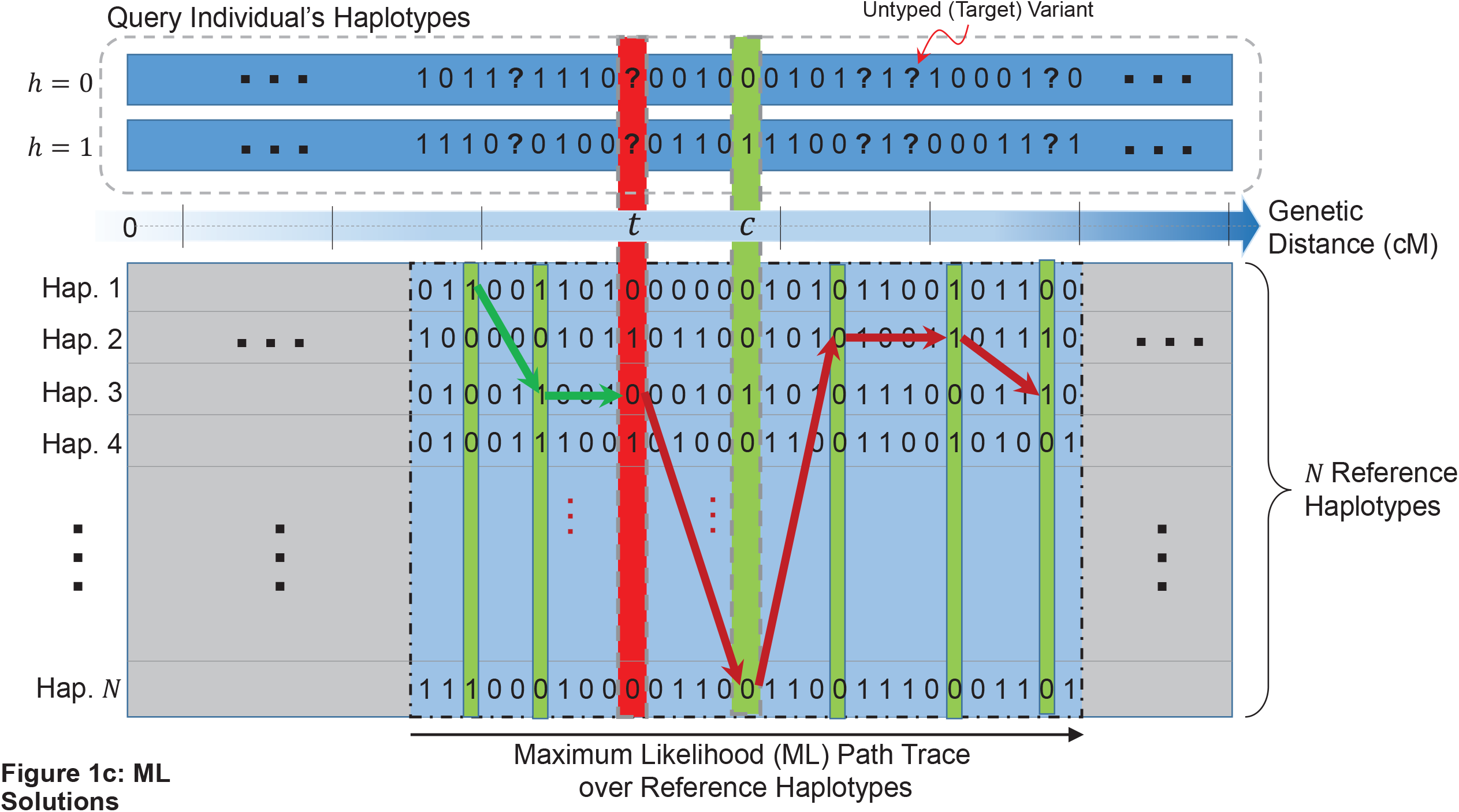
(a) Illustration of the local imputation setup. Query individual’s parental haplotype copies are shown in two rectangles in top, which are strings of {0,1}. 0 and 1 indicate a reference and alternate allele, respectively, for corresponding variants. The genetic distance (in centimorgans) are shown with the blue arrow and is used to track the center position of the window and the target untyped variant. The reference haplotypes are shown in the box below wherein each row corresponds to a haplotype. Given the local window of radius *l*_*win*_ the window is illustrated in the dashed rectangle whose center is shown at the genetic position *c* and for the target variant at position *t*. The typed variants are shown in green rectangles and the untyped target is shown in the red rectangle, whose alleles on the query haplotypes are shown with question marks. (b) Illustration of the forward and backward variables. For the 3^rd^ haplotypes at the untyped variant, the incoming paths (forward variable) are illustrated with green lines. The outgoing transitions are shown with red lines. The self-transitions are shown with heavier lines compared to the non-self transitions. (c) The ML (Viterbi) path is shown with the transitions along the haplotypes.

#### Li-Stephens Markov Model

Our evaluations use the Markov model defined by the standard Li-Stephens model^40^, where the haplotypes of each query individual are modeled as a “mosaic of the reference haplotypes” such that pieces of reference haplotypes (consecutive variant alleles on a haplotype) are concatenated to each other. This model describes a probability distribution on possible “paths” that pass over the reference haplotypes. In this model, the transitions between the haplotypes and errors on the haplotypes are probabilistic. In the simplest sense, the minimal number of haplotype transitions and allelic errors can be thought of as the most likely path that describes the query haplotype. The basic idea is to pin-down the tag variant alleles on the paths, and estimate the probability of alleles at the untyped variants:

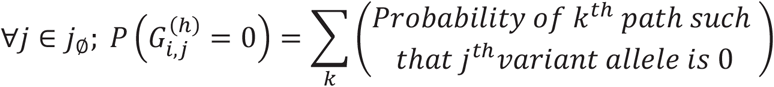

In this model, the haplotypes of the reference panel correspond to the states of the Li-Stephens Markov model. Each state (haplotype) emits an allele at a variant position *1* ≤ *j* ≤ *V*. In addition, the transitions between the states (i.e., the switches between haplotypes) at variant *j* are dependent only on the genetic distance between the variants at indices *j* and (*j* + 1). The genetic distance measures the probability of recombination taking place between these two variants. In the Markov model, recombination corresponds to a state-switch whereby the state (i.e., the haplotype) makes a transition to a new state. However, the recombinations occur as homologous chromosomes crossover in the course of meiosis. The rate of recombinations changes depending on the position on the genome, i.e., some parts of the genome are more likely to harbor recombinations than others. Thus, the prevalence of recombination events along the genome is quantified in terms of genetic distance that is measured in centimorgans (cM), a measure of recombination probability between two loci. Given two variants at indices (*j* − 1) and *j*, the probability of recombination is modeled as:

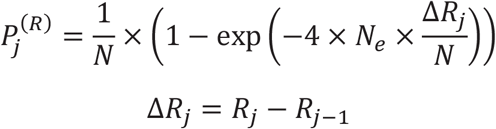

where 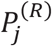 denotes the probability that there is a recombination event (i.e. Markov chain stays on the same state), Δ*R*_*j*_ denotes the genetic distance between variants at indices (*j* − 1) and *j*, and *N*_*e*_ denotes the effective population size. It is important to note that the probability of recombination depends only on the position of the variant and not the actual haplotype. This is widely used in HMM-based imputation methods to decrease computational costs. The probability of that a recombination does not take place can be computed from 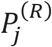:

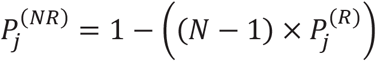

where all recombination events are accounted for and removed from 1 and 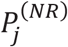 indicates that there is no recombination between variants at indices (*j* − 1) and *j*. From the above equation for 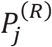, Increasing population size implies a higher probability of recombination, i.e., larger effective population size indicates more complex recombination patterns as the probability of switching between haplotypes (or states) increases. Given the query individual’s phased genotypes, 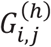 and the reference haplotype data, *H*_*i,j*_, the hidden Markov model is defined based on these equations using the transition and emission probabilities:

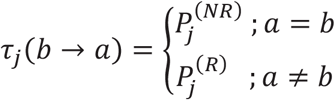

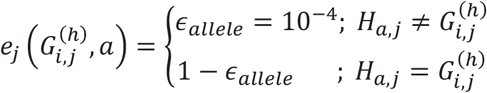

where *τ*_*j*_(*b* → *a*) denotes the transition probability from haplotype *b* to *a* at variant index *j* from the previous variant at index *(j* − 1) and 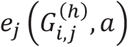 denotes the emission probability of the allele 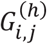 from the *a*^*th*^ haplotype. The emission probability depends on the alleles of the query individual; if the allele on the *a*^*th*^ haplotype matches query individual’s allele, a high emission probability is assigned, otherwise allele error probability, *ϵ*_*allele*_, is assigned as the emission probability.

Using the above equations and Li-Stephens Model, we use two approaches for inferring the haplotype states at every typed variant.

#### Inference of Posterior State (Haplotype) Probabilities

First approach the estimation of per-typed-variant estimate of posterior probabilities of each haplotype and assignment of the maximum *a-posteriori* (MAP) estimate of the alleles at the untyped variants. For this, we make use of the forward-backward algorithm^52^, which is a well-known dynamic programming algorithm that is used to efficiently compute the state probabilities at each step of the HMM as

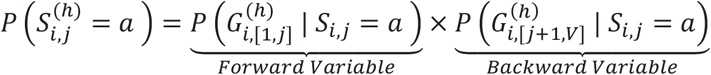

where 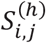 denotes the state (haplotype) of the HMM at variable index *j* for individual *i*’s parental copy 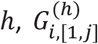 denotes the sequence of alleles for variants in [1, *j*] on individual *i*’s parental copy *h*, and 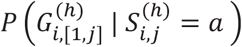 is the forward-variable and it denotes the probability of emitting the allele sequence 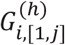 given that HMM is at state *a* at the variant position *j*. Backward-variable is similarly defined for the rest of the allele sequence that is backward of *j*^*th*^ variant, i.e., 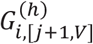. The forward and backward variables are computed using efficient recursion relations (See Methods). After the posterior probability at each variant position *j* and for each state *a* is computed, we can estimate the posterior probability of each allele at each untyped position:

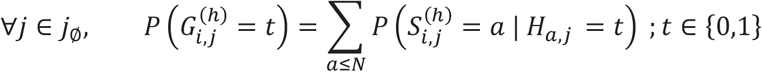

where the untyped variant allele *t*’s probability is estimated by marginalizing over the states *a* for which the corresponding haplotype has an allele *t*. As we describe below, we evaluate 2 different approaches for marginalizing over the haplotypes.

#### Maximum-Likelihood Mosaic-Haplotype (Viterbi)

Vicinity-based HMMs focus only on the vicinity of untyped variants. The main idea is to select a vicinity for imputation of a target variant.

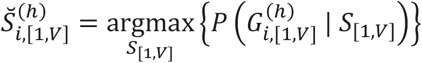

where 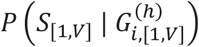 denotes the likelihood of the state sequence *S*_[1,*V*]_ given the allele sequence of all variants in [1, *V*] for *i*^*th*^ individuals. 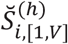 denotes the maximum-likelihood (ML) state sequence for *i*^*th*^ individual’s haplotype *h*. This state sequence represents the most likely mosaic haplotype that gives rise to the variant alleles 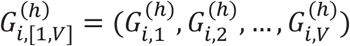. This state sequence can be inferred using a dynamic programming algorithm, namely the Viterbi algorithm^53^ that efficiently identifies the maximum-likelihood state sequence similar to the forward algorithm.

After the ML state sequence is computed using the Viterbi algorithm, we can assign the alleles on the haplotypes that correspond to the ML state sequence at every variant:

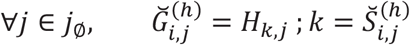

where 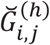 denotes the ML allele for the untyped variant *j* that is assigned to individual *i*’s haplotype *h*. The Viterbi algorithm does not immediately assign a score for the ML allele. We aggregate the vicinity information to assign a score for the imputed ML allele.

#### Locality Parameters of Imputation

We evaluate the effect of changing parameters on the accuracy of genotype imputation. The forward-backward and Viterbi-based imputation algorithms sequentially analyze the variants while keeping track of the scores and probabilities for each state. They can be performed using all of the variants on each chromosome as the LD information is confined generally to individual chromosomes and inter-chromosomal LD information, while detectable, are very weak^54^. These are out-of-scope of the imputation methods that we evaluate. Using whole chromosomes in imputation enables the algorithm to integrate the linkage information from all positions on the chromosomes. On the other hand, the linkage information tends to decrease quickly while imputing an untyped variant, e.g., the identity-by-descent segment length (length of conserved haplotypes) decreases quickly among generations (25 generations separation have on average 2cM conservation^36^). This information can be integrated into MAP and ML-based imputation by a sliding-window where the variants outside a local window are not used for imputation. This can help decrease the computational requirements. For example, BEAGLE uses a large sliding window (length 30 cMs) and merges the consecutive windows to infer the forward and backward variables. In our study, we run forward-backward and Viterbi algorithms solely on the local windows around the untyped variants and use these “local-HMMs” to impute the untyped variants. For instance, if we are using a local window of length 0.5 cMs, the ML state sequence is computed only for the local vicinity of the is assigned as:

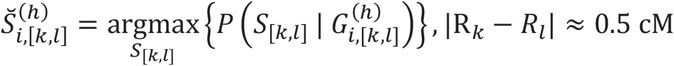

where the ML state sequence, 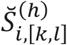, is confined to the variant indices [*k, l*] whose genetic distance is approximately 0.5 cMs. The forward-backward computations are similarly confined to the local windows based on genetic distance cutoffs.

We test different local window lengths and evaluate the impact of window length on the allele imputation accuracy. We utilize a sliding window with lengths from 0.1 cM upto 1 cMs and compute the imputation accuracy (See Metrics). Another important factor is the positioning of the untyped target variant within the local window. It is expected that the LD information can be integrated more accurately if the untyped variant is centered around the local window. It is, however, not clear to what extent “target-to-center distance” affects the imputation accuracy. For each untyped target variant, we first identified the typed variants that will be used for imputation that satisfies the local window length and target-to-center distance criteria using ML and MAP approaches with the selected population size and allelic probability assignment procedure.

#### Evaluation Setup and Metrics

We use genotype data from the 1000 genomes Project’s Phase 3^55^. We focus on the variants on chromosomes 19, 20, and 22 for extensive evaluation and exclude the multi-allelic SNVs and indels. Among these data, chromosome 22 is used to evaluate different parameter combinations. To decrease computational requirements with the parameter combinations, we focused on the region chr22:25,000,000-35,000,000. In the evaluations, we randomly selected 1000 individuals as the phased reference panel and 200 individuals (with known genotypes) for estimating evaluation. After we evaluated the parameters, we selected the optimal parameter set and validated the imputation on chromosomes 19 and 20. To define the typed (tag) variants, we extracted the positions of the variants that are genotyped on the Illumina Duo 1M genotyping array platform^56^. This enables us to perform evaluations on a realistic test case as the Illumina’s array is used in several large-scale projects including the HAPMAP project^56^. We used all the variants that map to the positions that overlap with Illumina Duo platform as the typed variants and the remaining variants are assigned as untyped variants that are imputed. After extracting the variants, we phased the genotypes using EAGLE2^51^. We did not rely on the phasing that is provided by 1000 Genomes because it may artificially increase the accuracy of the imputation accuracy for the local HMM approach in comparison to established methods. The phased typed variants are input to LoHaMMer with different parameters for imputation. After the untyped variants are imputed, we computed the (1) “Genotype R^2^”^57^, (2) Genotype concordance (all and non-reference genotypes), and (3) Precision-Recall curves based on the imputed probabilities. We compare the implemented locality-based HMMs with BEAGLE, which is used as the baseline method for imputation. The imputation accuracy is classified among variants with respect to the range of minor allele frequency (MAF) and with respect to the chromosomal position.

### Evaluation of Imputation Accuracy with Changing Locality Parameters

For assessing the imputation accuracy with changing parameters, the variants are classified into “common” (MAF>0.05) and “uncommon” (MAF<0.05) variants. We tested the impact of the 4 different parameters, (*l*_*w*_, *N*_*e*_, *l*_*c*2*t*_, *n*_*tag*_) that describe the local HMM window around the untyped (target) variants. Rather than computing all parameter combinations, we selected a range for each parameter and we evaluated the impact of one parameter while keeping others constant. We used (*l*_*w*_, *N*_*e*_, *l*_*c*2*t*_, *n*_*tag*_) = (0.3,10^4^, 0.05,10000) as the default parameter values.

#### Local Window Length (l_w_)

We first evaluated the impact of local window length on the imputation accuracy. We used local window lengths of *l*_*w*_ ∈ {0.02, 0.05, 0.1, 0.2, 0.3, 0.4, 0.5, 1} cM. Figures 2c and 2d show the genotype R^2^ distribution and non-reference genotype concordance of the common variants on chromosome 22 with the changing local window length. The imputation accuracy of the 3 methods, MAP, ML, and BEAGLE (baseline) are reported. As expected, we observed that the accuracy increases with increasing window lengths. For window lengths above 0.3 cMs, we observed that there is less than 0.05% increase (91% vs 90.5%) in the R2 accuracy and similarly, 0.5% increase in the non-reference genotype concordance when MAP and BEAGLE are compared. For the uncommon variants, the minimum window length greater than 0.5 cMs exhibits very similar behavior as BEAGLE (0.5% decrease). Figure 2e shows the sensitivity and precision (i.e., positive predictive value – PPV) of the imputed genotypes with respect to changing impute genotype probability. For each window length, a curve is plotted. As the window length increases, both the sensitivity and precision increases. BEAGLE exhibits the optimal baseline performance. Above 0.3 cMs, the curves are very close to each other. These results indicate that *l*_*w*_ > 0.3 cM is the minimum window length with comparable accuracy as BEAGLE. For uncommon variants, we observed that most of the *R*^2^ and concordance are at the high or low accuracy regimes (Fig. 3a,b) for both BEAGLE and LoHaMMer. The PR curves demonstrate a fairly steady pattern of change in the accuracy (Fig. 3c,d).

**Figure 2.**
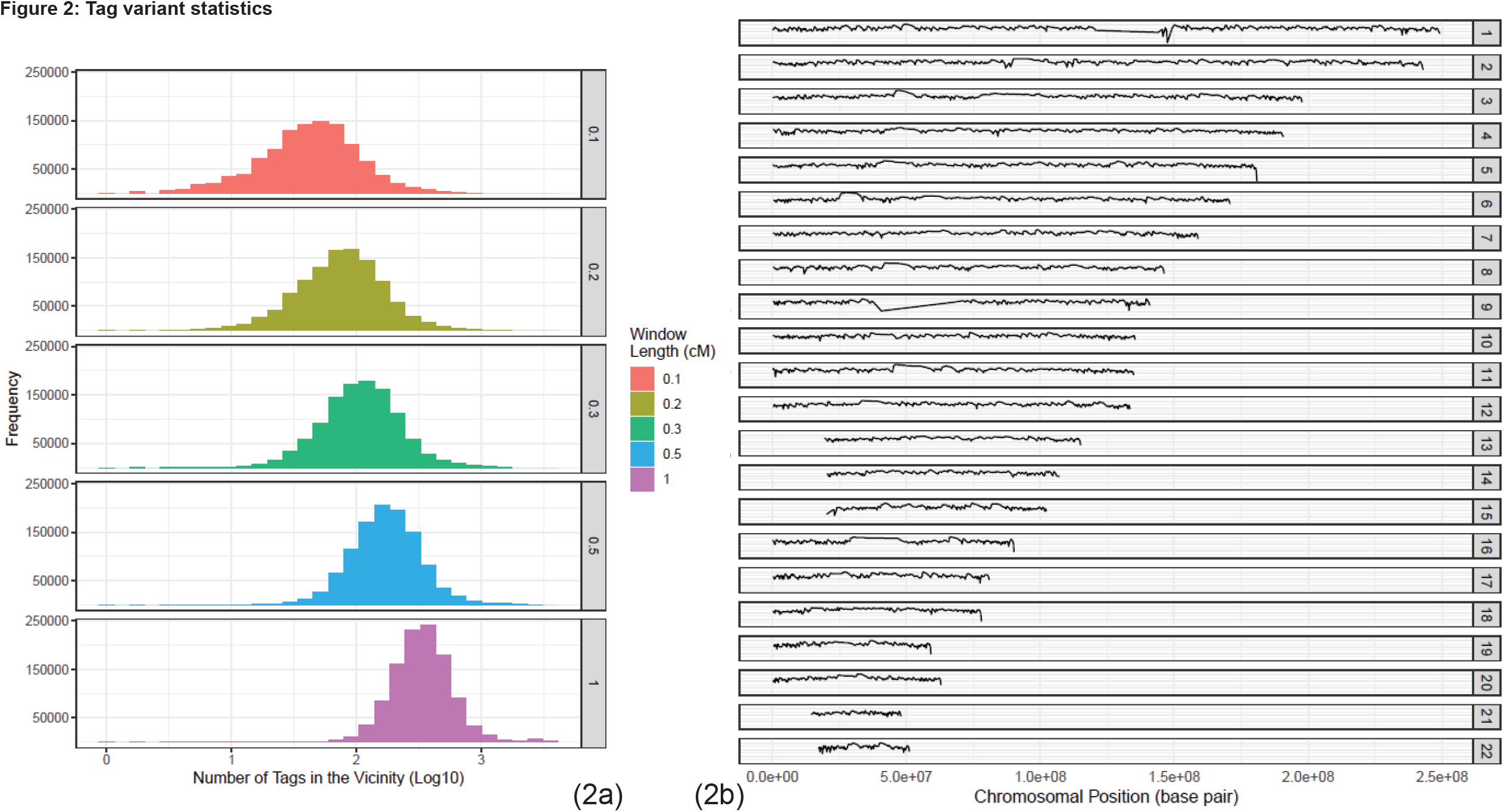
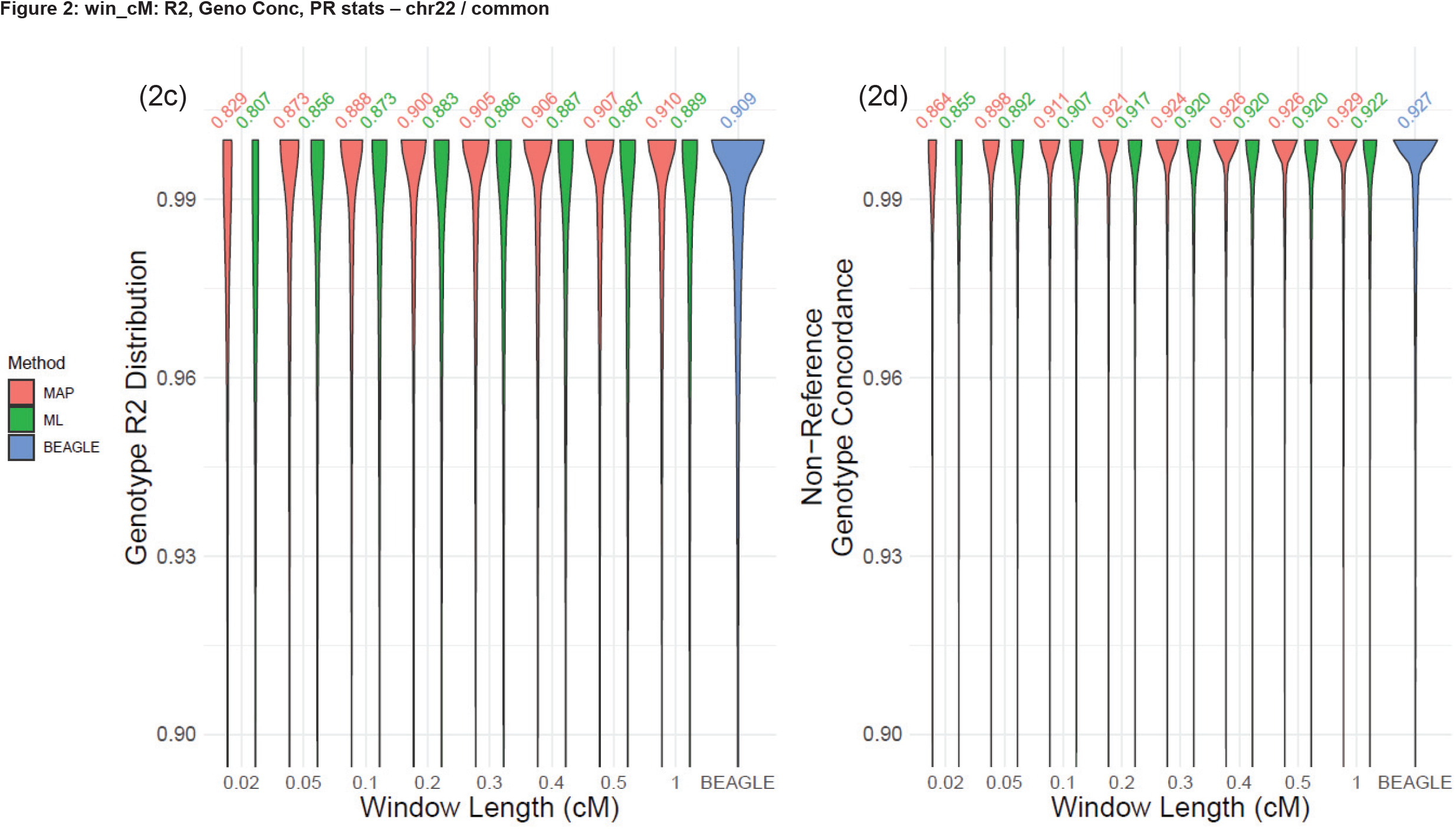
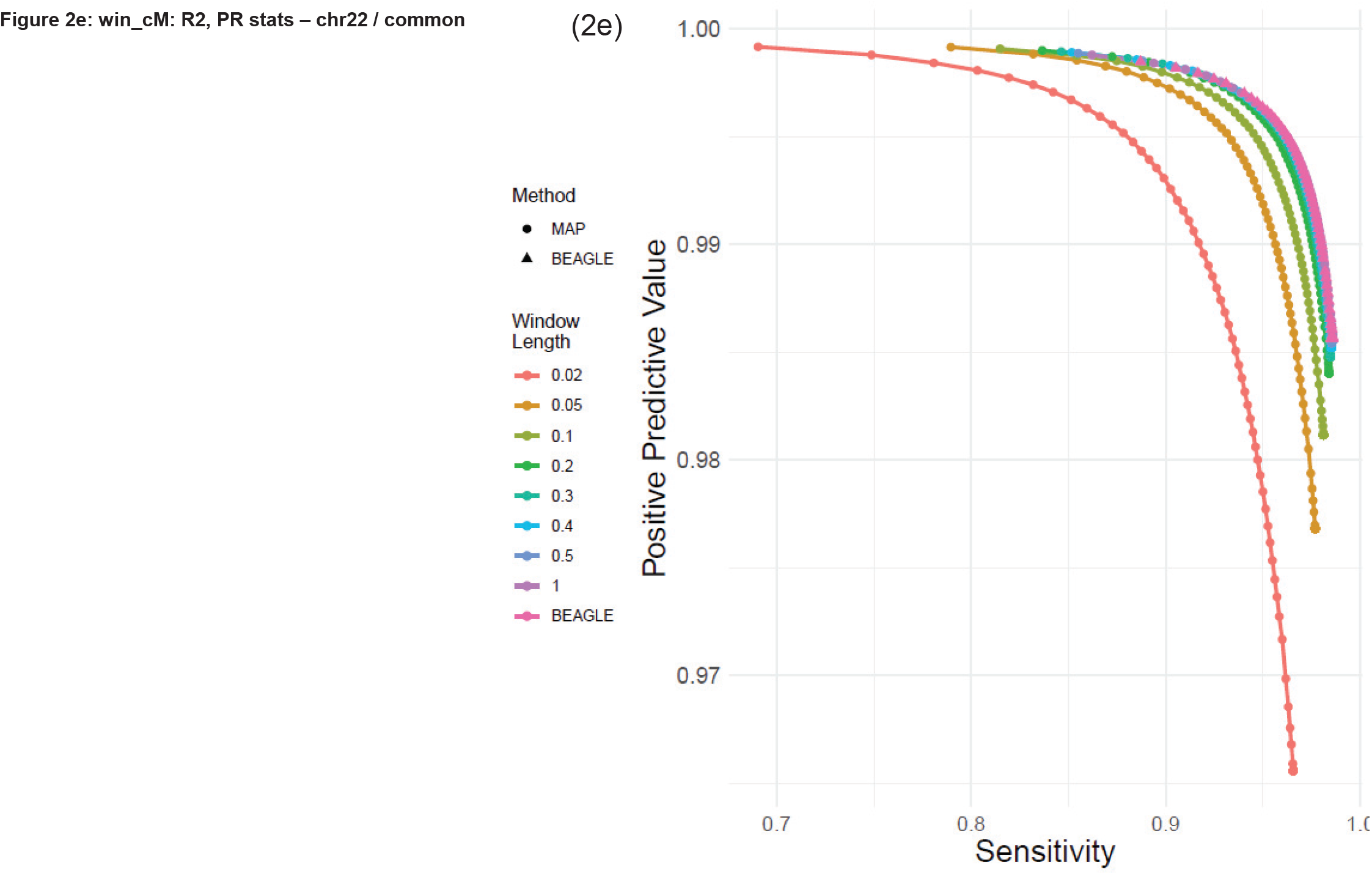
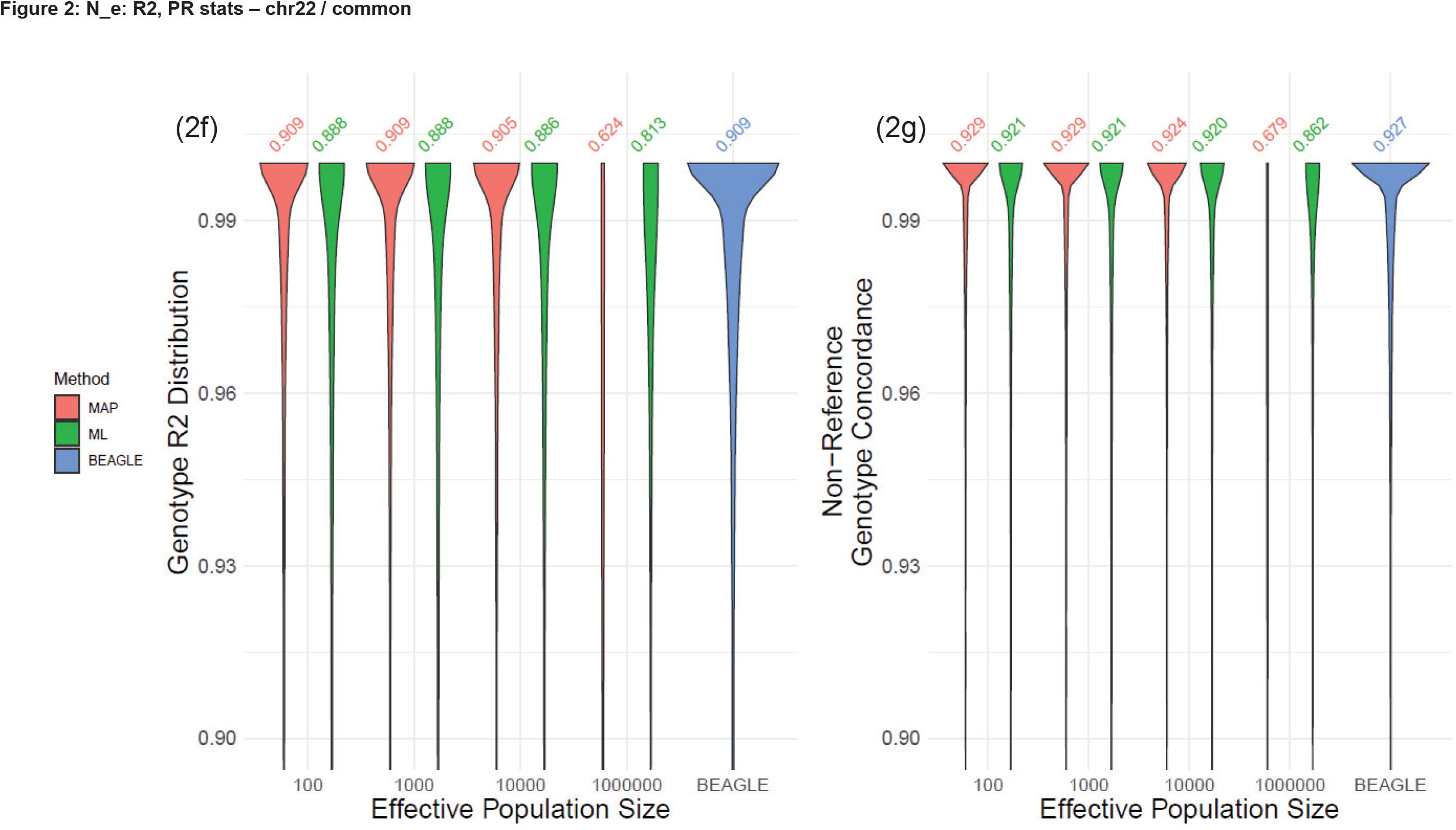
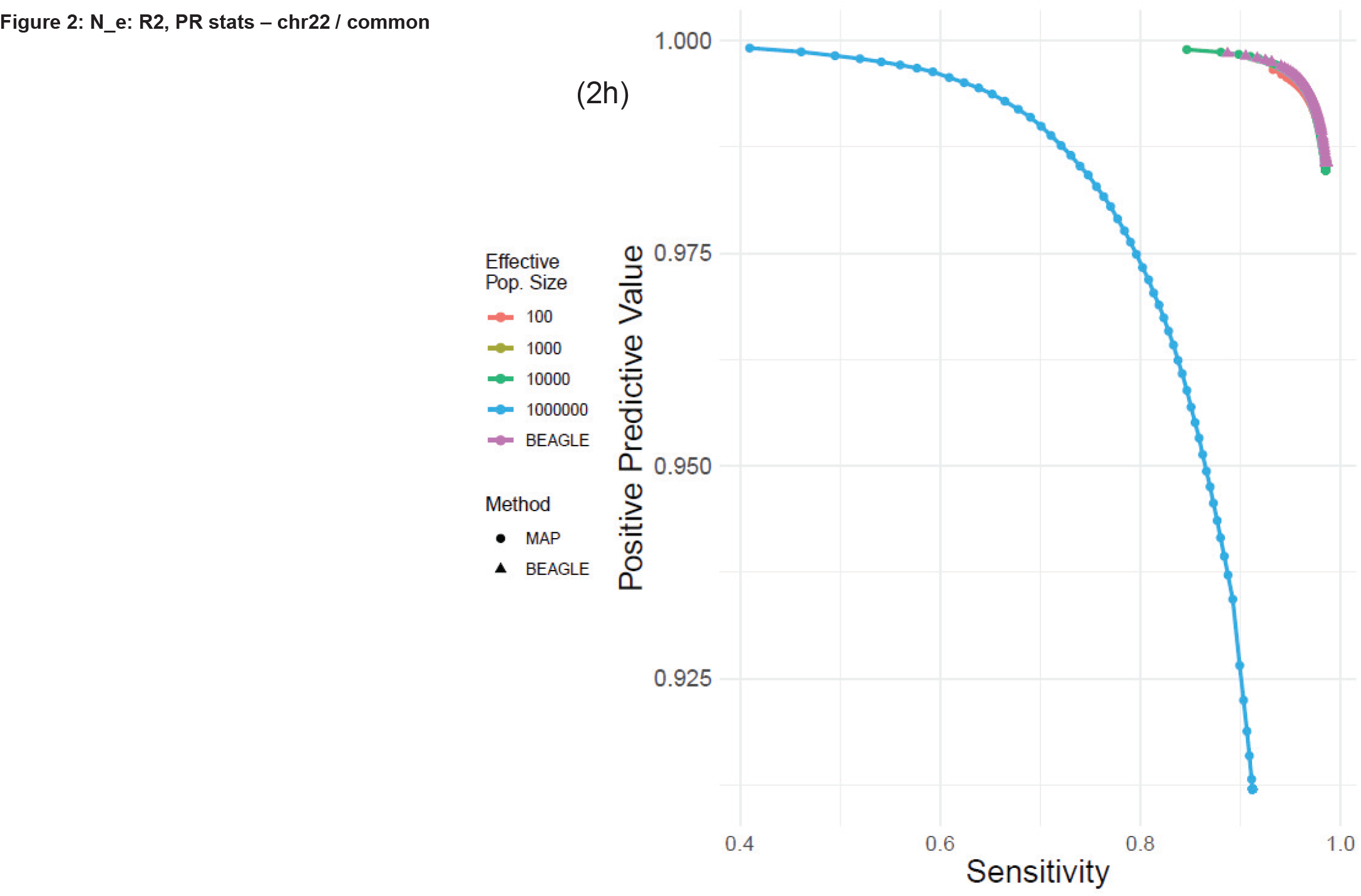
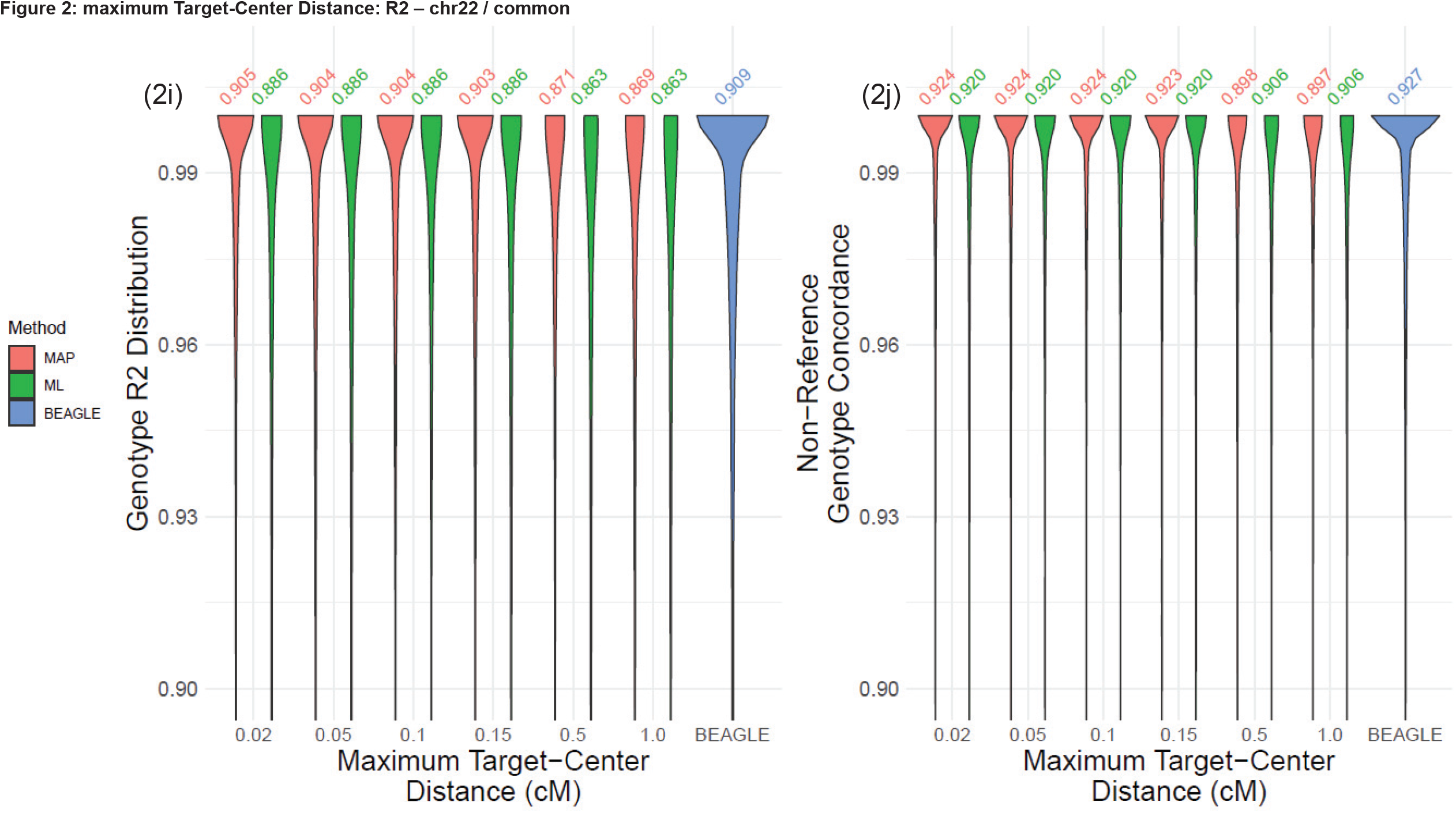
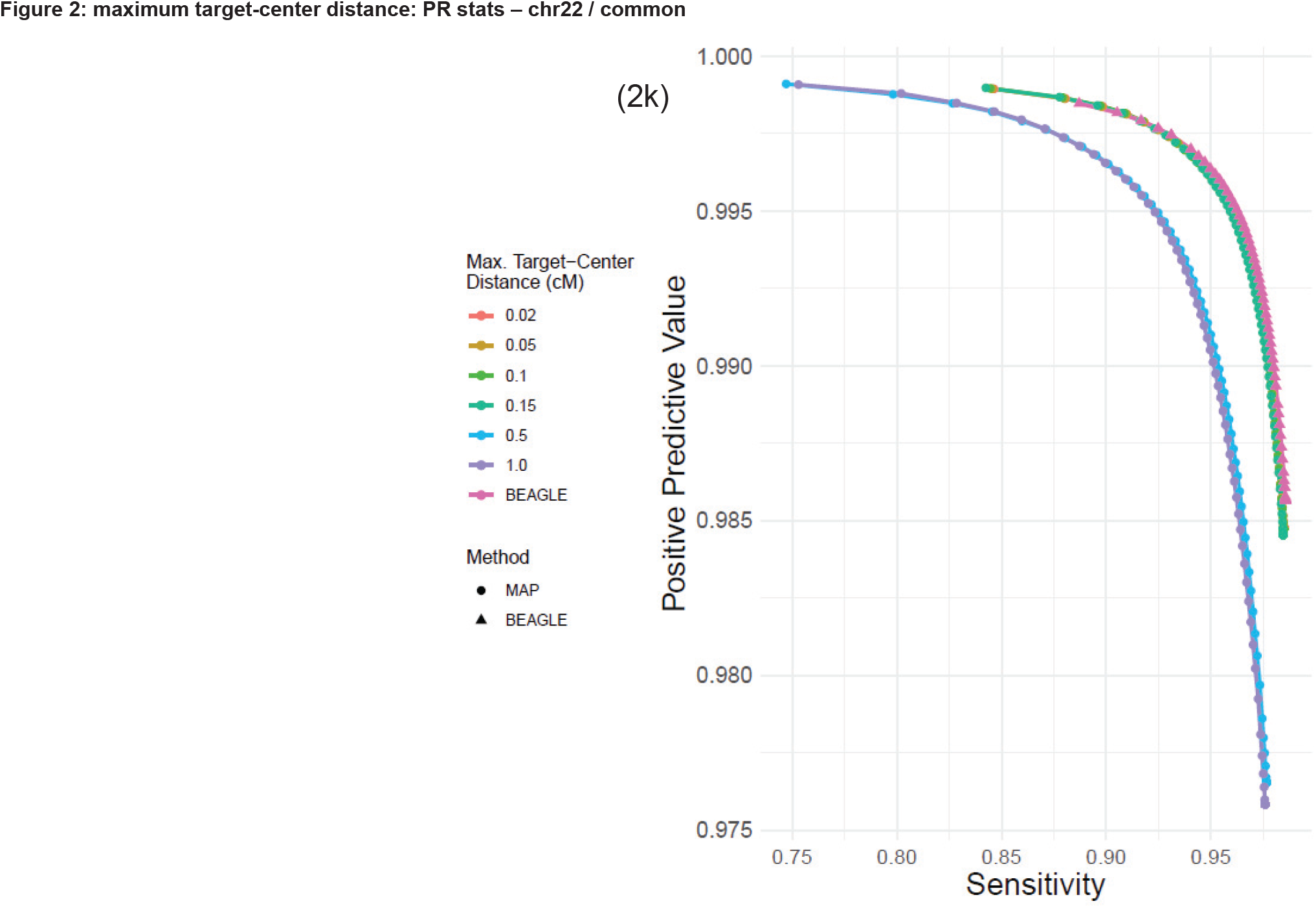
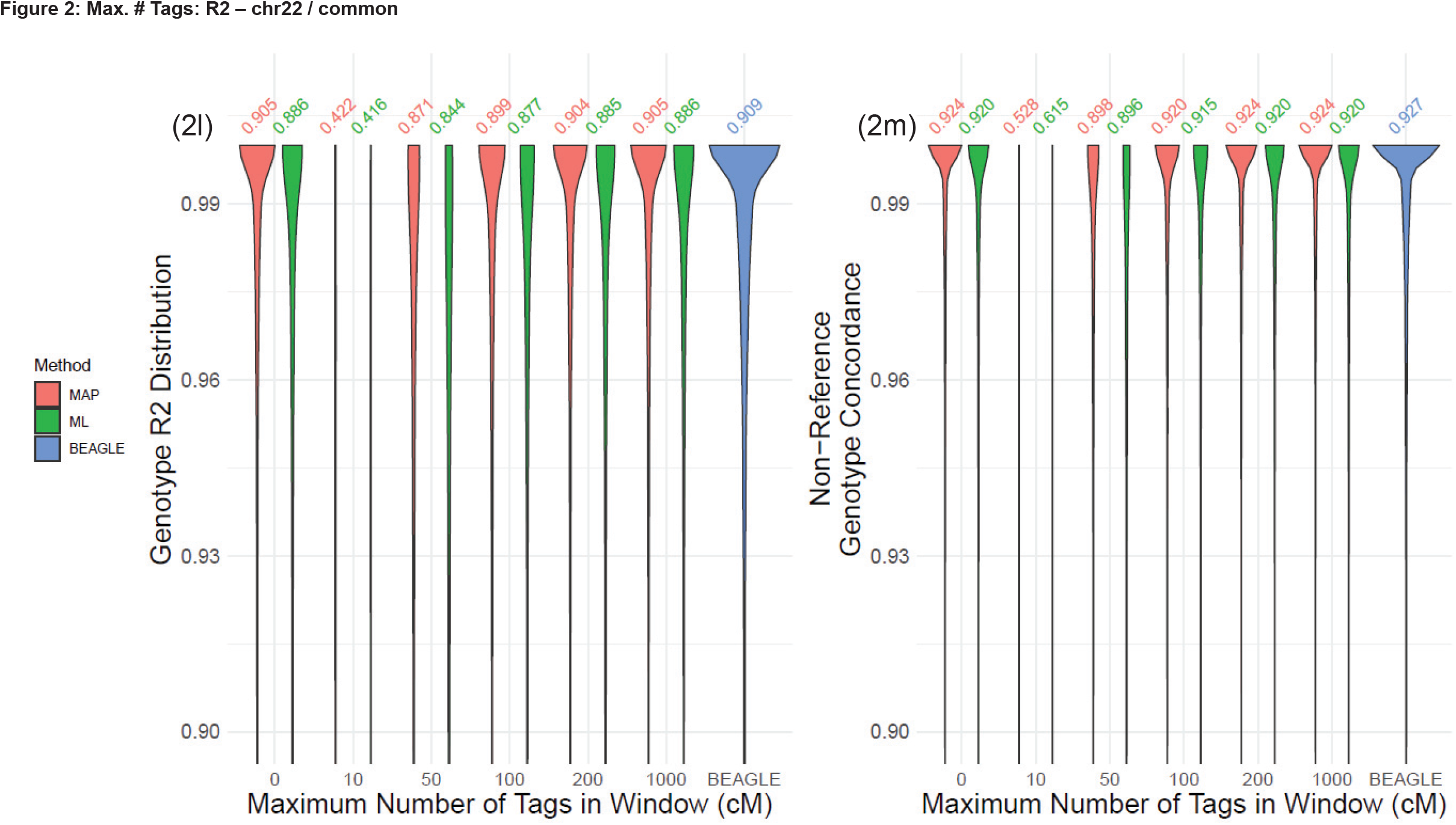
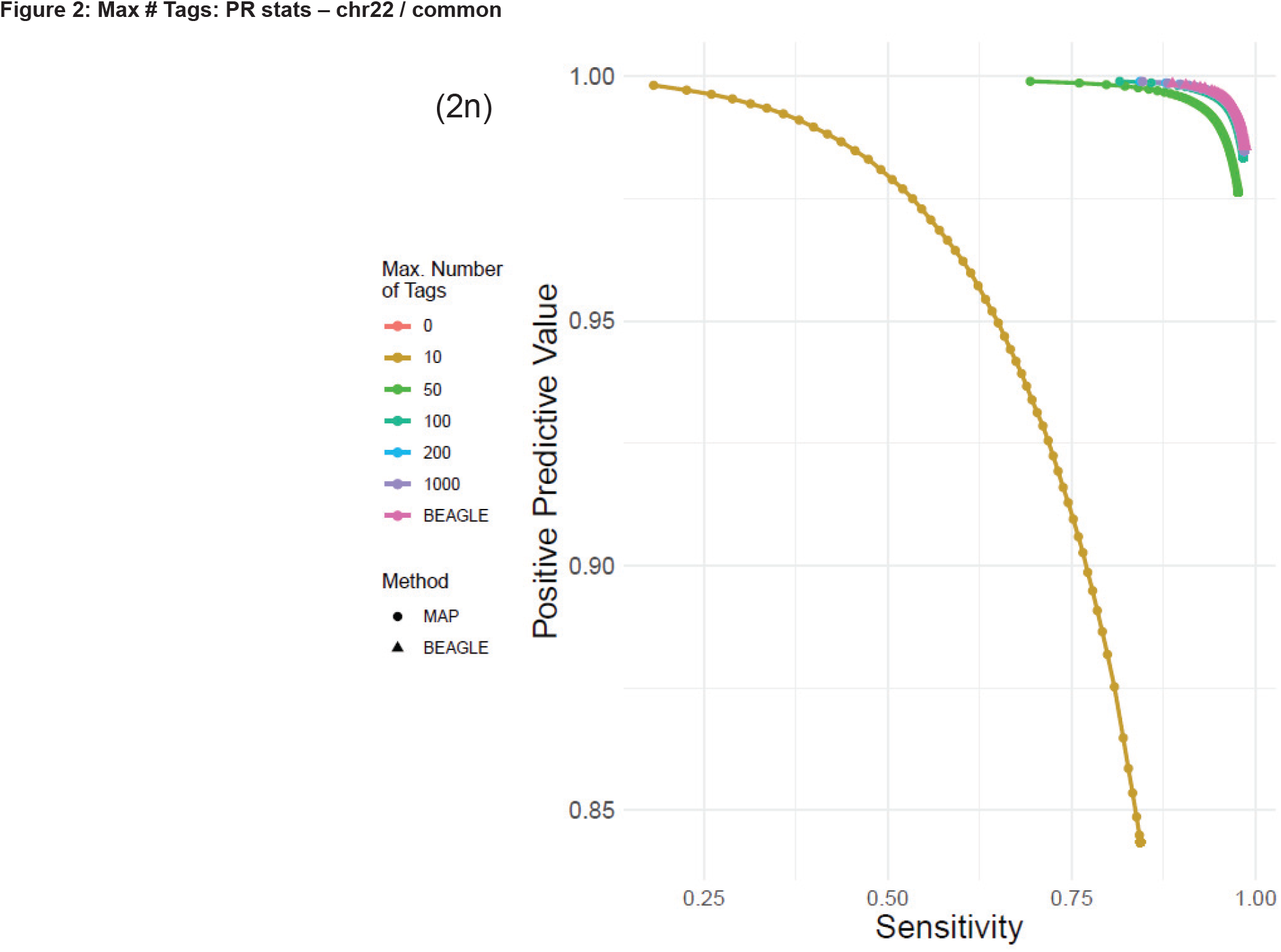
(a) The distribution of typed variant number for different window lengths for common untyped variants. (b) The number of tag variants for 1 cM over the chromosomes, each row is a chromosome and x-axis shows the chromosomal position. (c,d) Distribution of genotype *R*^2^ and genotype concordance for changing locality window length (*l*_*win*_). (e) The PR-curve for changing *l*_*win*_. (f,g) Distribution of genotype *R*^2^ and genotype concordance for changing population size (*N*_*e*_) (h) PR-curve for changing *N*_*e*_. (i,j) Distribution of genotype *R*^2^ and genotype concordance for changing target-center distance, *l*_*c*2*t*_. (k) PR-curve for changing *l*_*c*2*t*_. (l,m) Distribution of genotype *R*^2^ and genotype concordance for different number of targets. (n) PR-curve for different number of targets.

**Figure 3.**
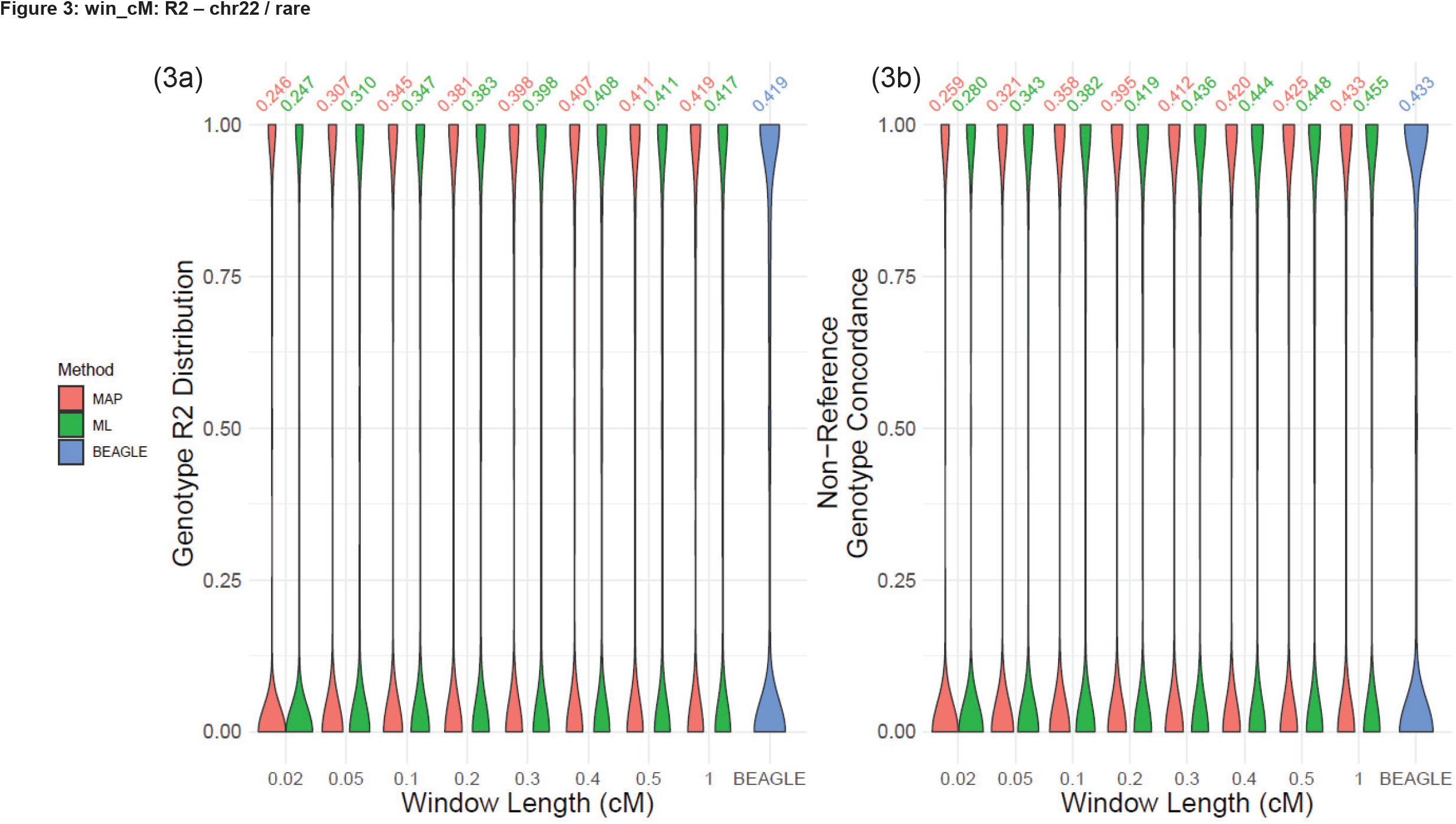
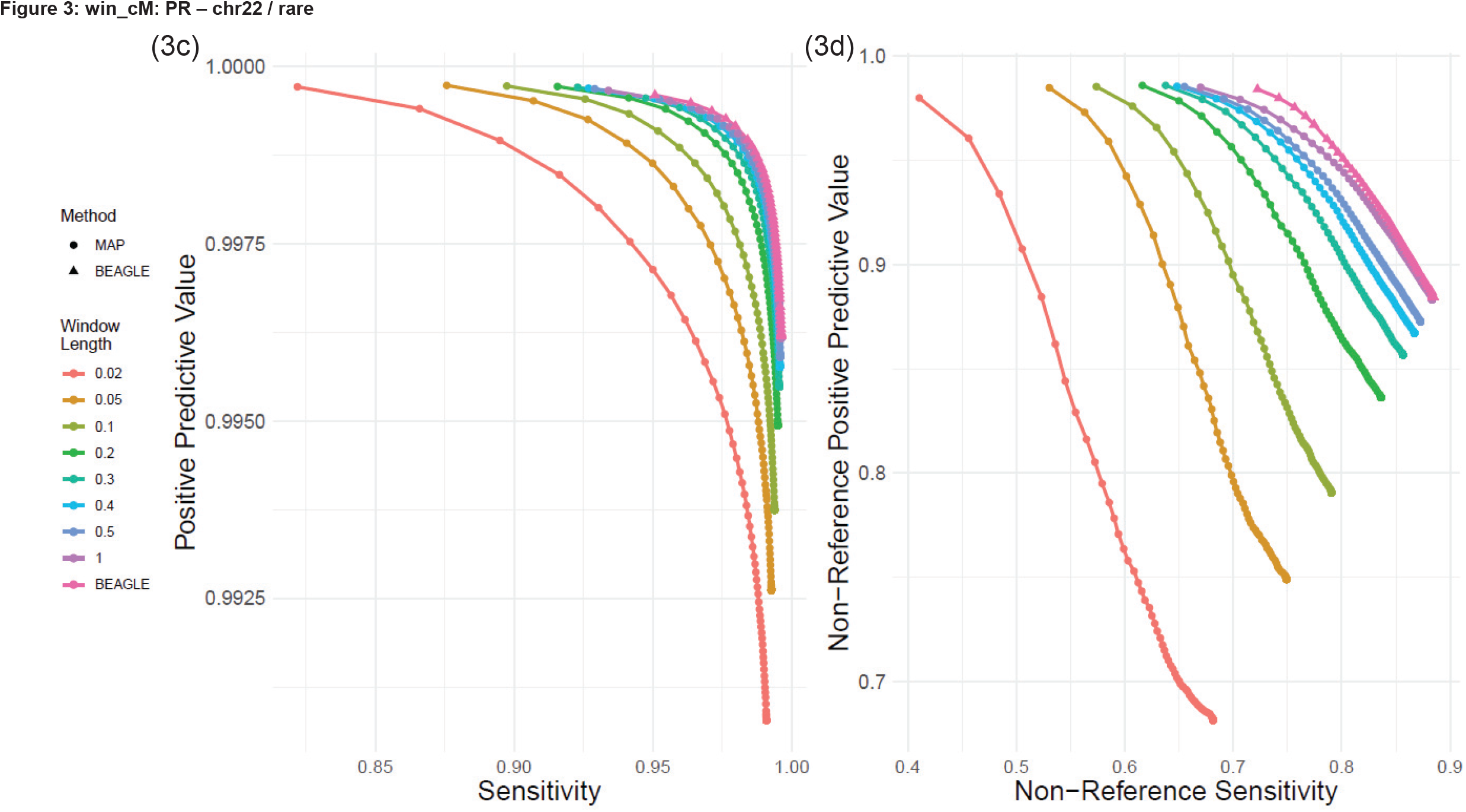
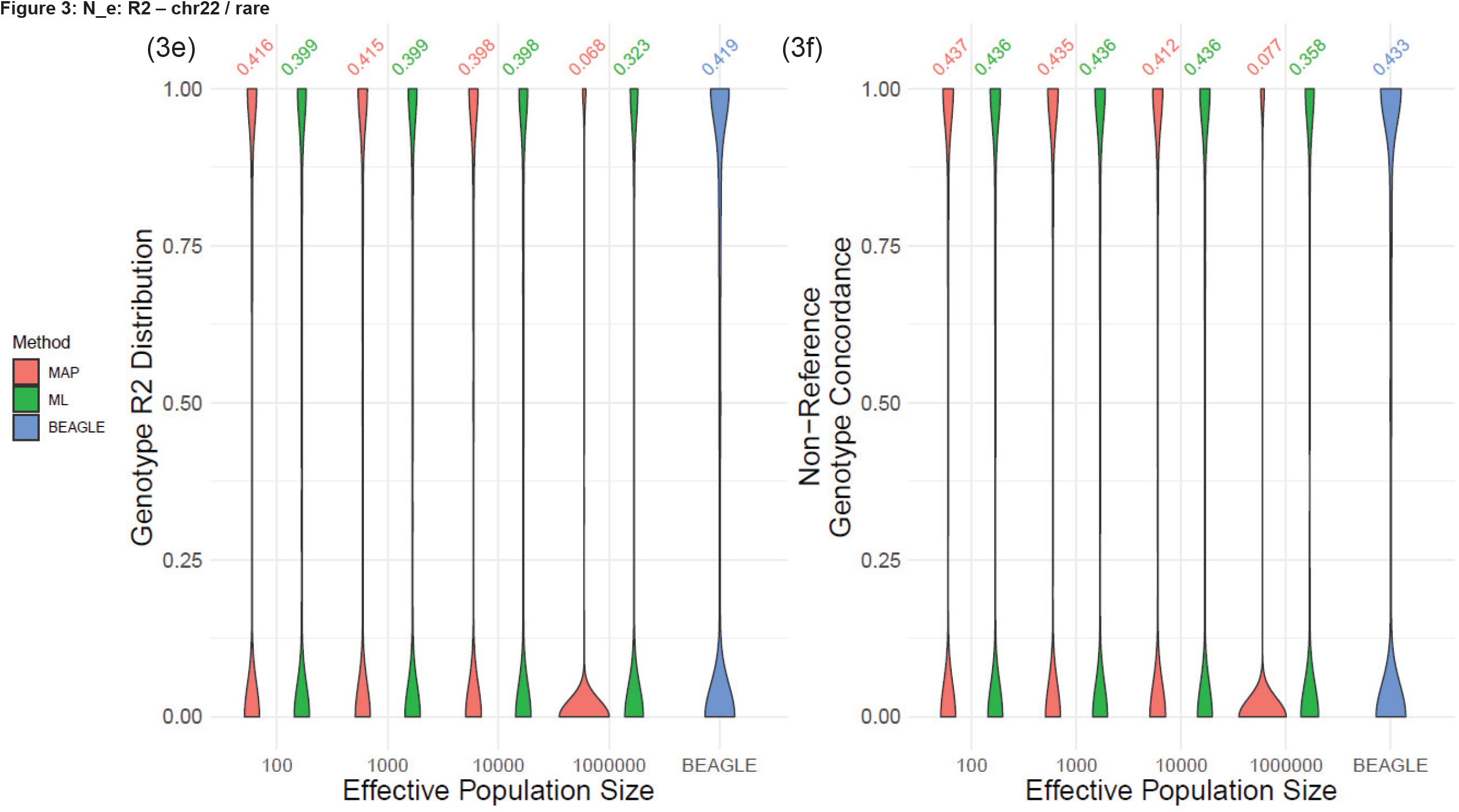
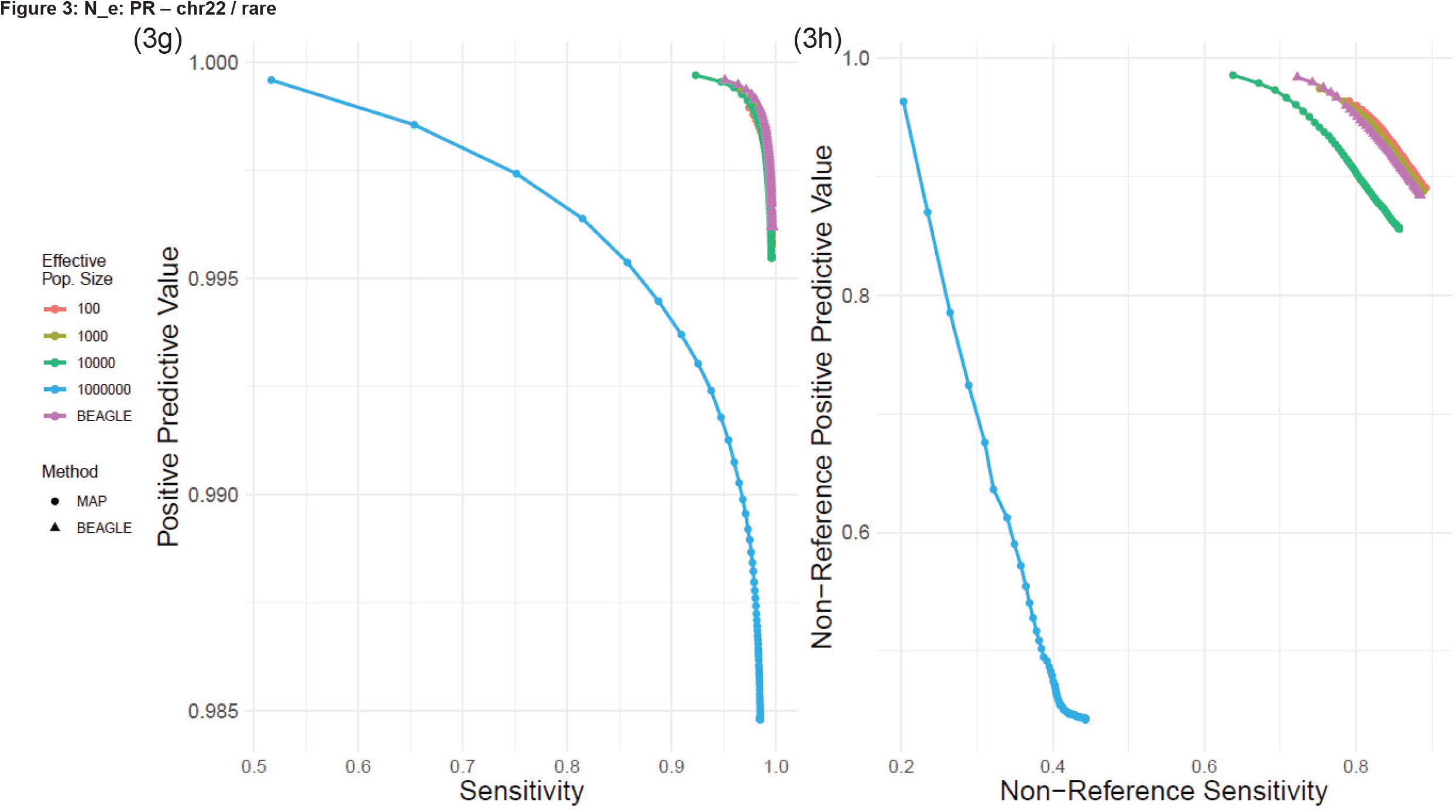
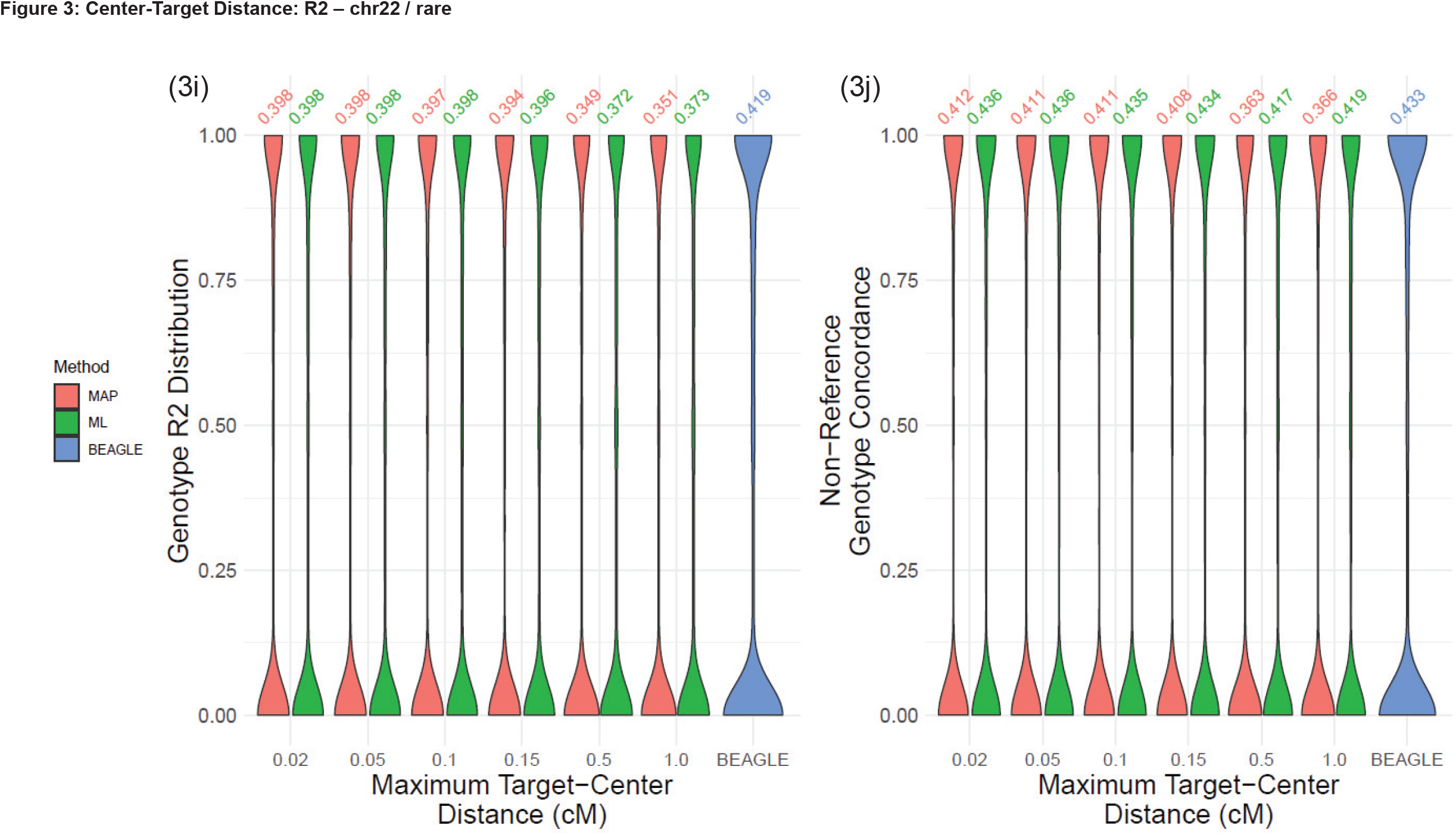
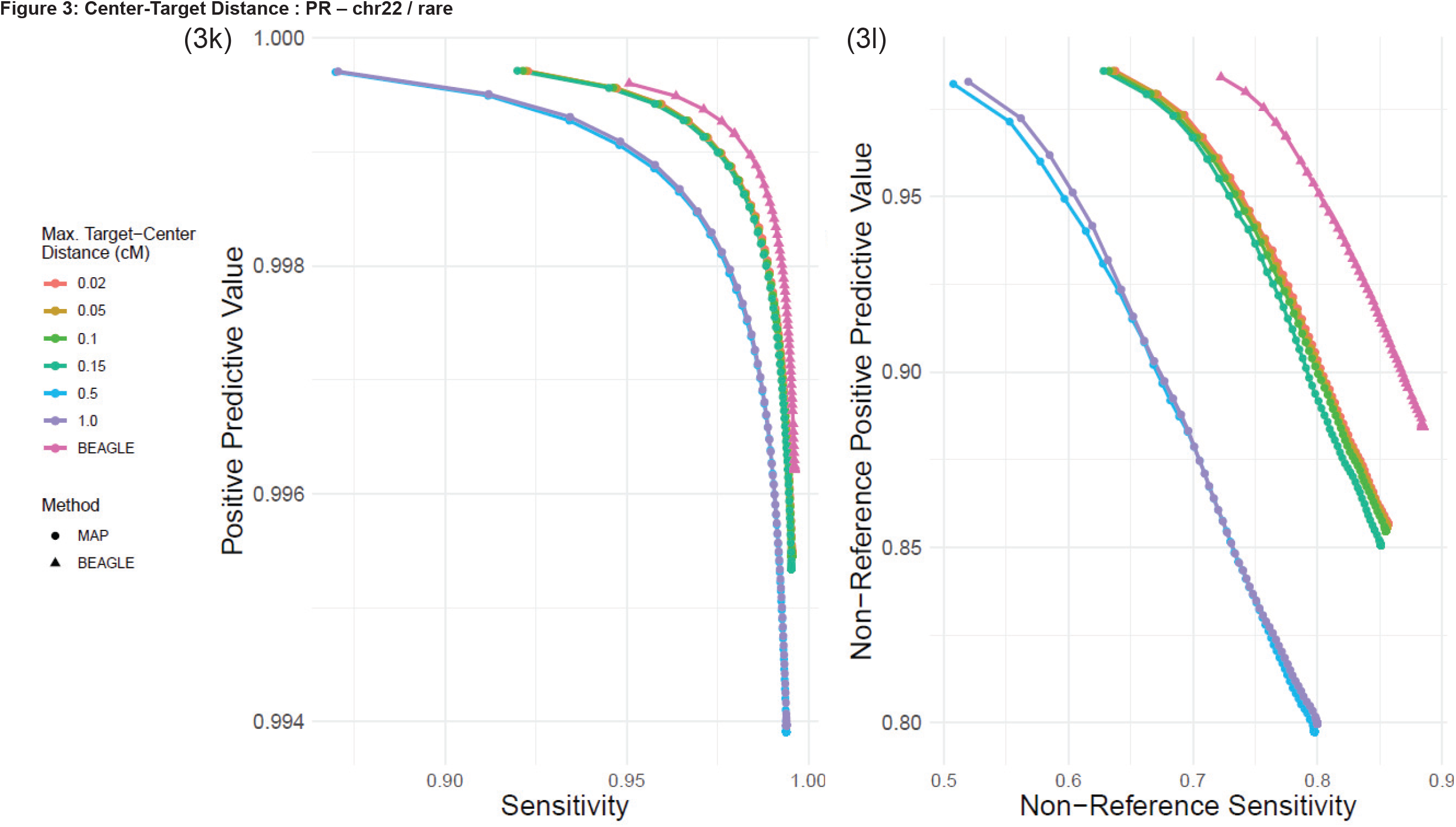
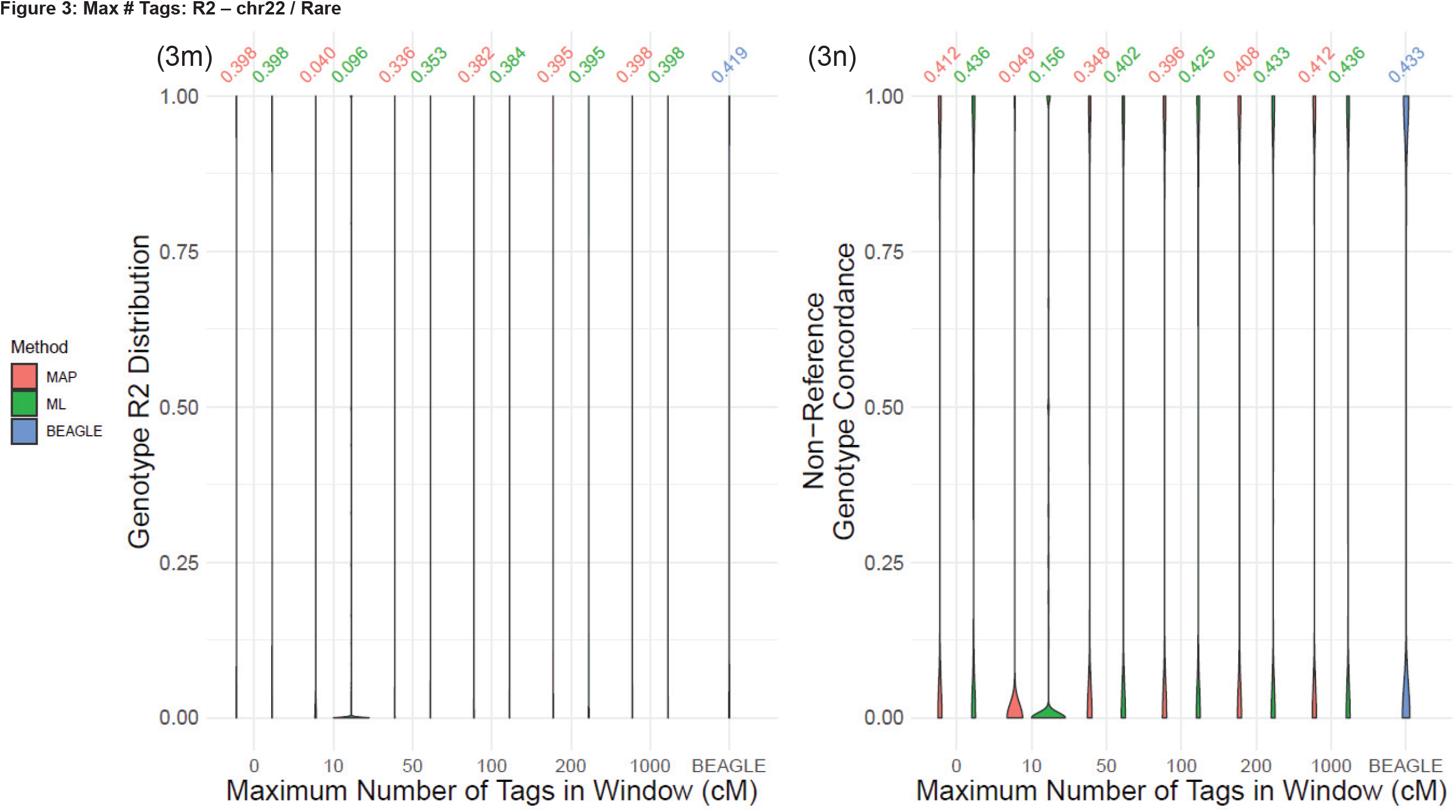
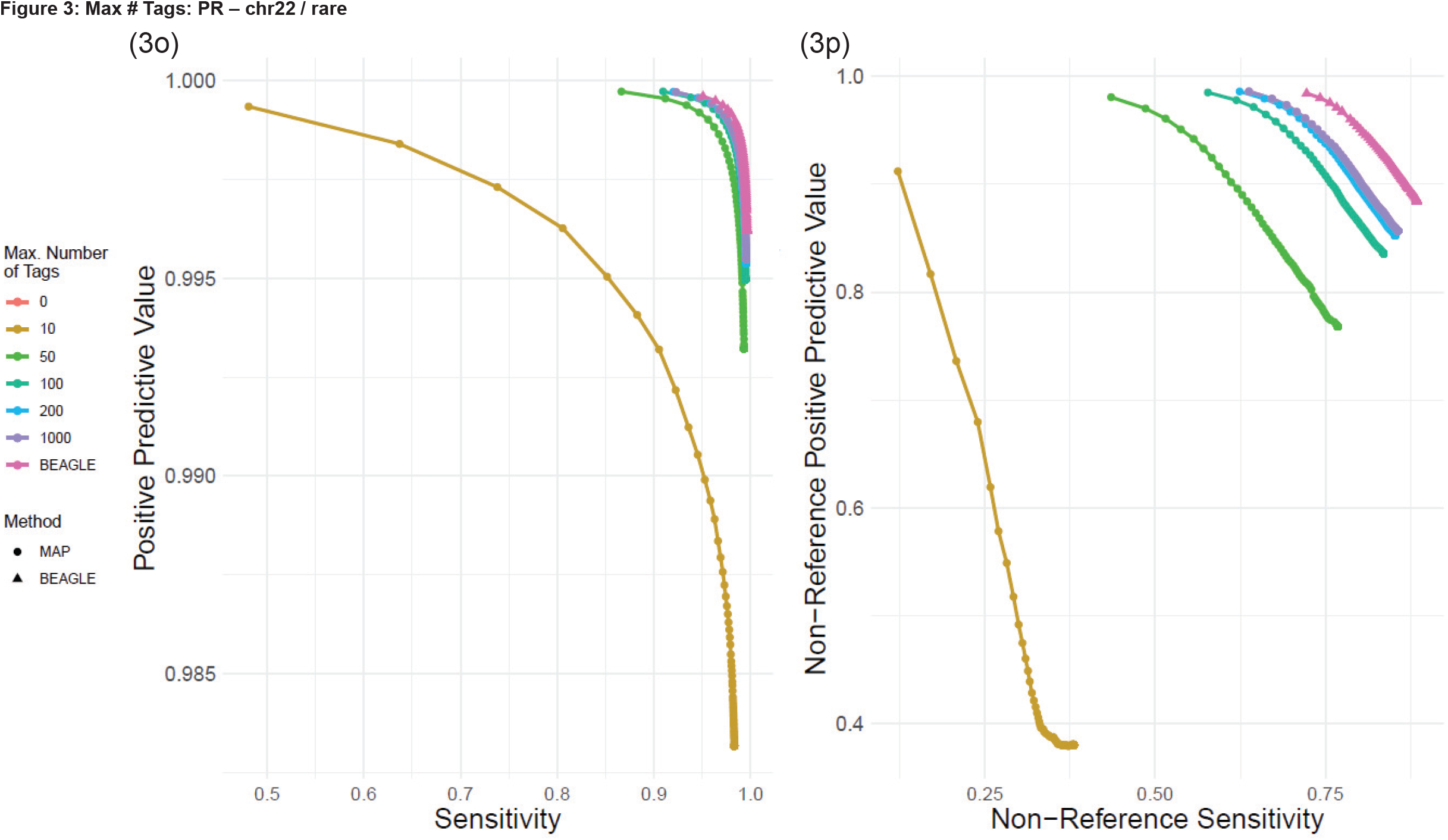
(a,b) Distribution of genotype *R*^2^ and genotype concordance for changing locality window length (*l*_*win*_). (c,d) The PR-curve (all and non-reference) for changing *l*_*win*_. (e, f) Distribution of genotype *R*^2^ and genotype concordance for changing population size (*N*_*e*_) (g, h) PR-curves for changing *N*_*e*_. (i, j) Distribution of genotype *R*^2^ and genotype concordance for changing target-center distance, *l*_*c*2*t*_. (k, l) PR-curves for changing *l*_*c*2*t*_. (m, n) Distribution of genotype *R*^2^ and genotype concordance for different number of targets. (o, p) PR-curve for different number of targets.

#### Effective Population Size (N_e_)

We analyzed how effective population size impacts prediction accuracy. We tested effective population sizes using *N*_*e*_ ∈ {10^2^, 10^3^, 10^4^, 10^6^}. Figures 2f and 2g show the accuracy and sensitivity-precision curves, respectively. We observed that *N*_*e*_ ≈ 10^3^ provides the best performing population size. Also, increasing *N*_*e*_ too high renders the transition probabilities uniform among self-transitions and recombination events causing too many recombinations. In turn, the model tends to overfit into any haplotype data without too much discriminative power since the model allows a large number of recombinations without much penalty. This causes the low performance with increasing *N*_*e*_. For uncommon variants, the impact of population size is very distinctive, *N*_*e*_ ≈ 10^3^ provides substantially higher imputation accuracy compared to other higher parameters (Fig 2h). The increase in *N*_*e*_ causes a sharp decline in imputation accuracy. For the uncommon variants, while a similar trend was observed, we also found out the ML method performs better for large values of *N*_*e*_. This highlights that ML-based imputation can provide more robust imputation results for certain parameters (Fig. 3e-h).

#### Target-Center Distance (*l*_c2t_)

The positioning of the target variant, *l*_*c*2*t*_ (target-center distance), in the imputation window is another parameter that can impact imputation accuracy (Figure 1a). We tested the imputation of accuracy with increasing target-center distance values, *l*_*c*2*t*_ ∈ {0.02, 0.05, 0.1, 0.15} cM. We use genetic distance as the measurement of unit for these parameters since it is the most natural choice. Imputation accuracy is shown for different center-target distance values indicating a visible impact of target-center distance (Fig 2i, 2j, 2k). We observed that the imputation accuracy decreases as *l*_*c*2*t*_ increases. This expected as smaller *l*_*c*2*t*_ implies that the haplotype evidence from the two sides of the target variant is optimized. Below *l*_*c*2*t*_ < 0.1 cM, we observed that the local window-based imputation provides comparable accuracy. For uncommon variants, we observed similar pattern in the accuracy (Figure 3i-l). As with other parameters, the rare variant are more sensitive to changes in target-center distance.

#### Maximum Number of Tag (Typed) Variants in Window (n_tag_)

The last set of parameters we tested are the number of tag (i.e., typed) variants that are used for imputation. For this, we subsampled the tags in each window such that the number of tag variants is bounded by the maximum number of tag variants. For this, we evaluated the impact of changing *n*_*tag*_ ∈ {10, 50, 100, 200, 1000}. The typed variants in the windows that harbor less than *n*_*tag*_ tag variants are used as they are. Figures 2k and 2m show the average *R*^2^ and non-reference genotype concordance. For *n*_*tag*_ greater than 100 variants, we observed that the accuracy levels out with a slight increase, both for common. For rare variants, we observed that the accuracy flattens out around *n*_*tag*_ = 200. Similar conclusions can be seen from the sensitivity-PPV curve (Fig. 2n). For uncommon variants, we observed a similar trend (Fig. 3m-p).

Overall, these results demonstrate that the local-HMM computation can provide imputation accuracy comparable to the baseline methods, especially for the common variants.

### Accuracy on Chromosomes 19 and 20

In order to validate and compare the local HMM parameter accuracy on a different positions, we tested the parameters for the variants on chromosomes 19 and 20. We extracted the typed variants on the Illumina Duo array platform on chromosomes 19 and 20. After this, we extracted 24,333 of 27,403 typed variants on chromosome 19 and 26,405 of the 28,319 typed variants on chromosome 20. The remaining variants (768,292 variants on chr19 and 742,370 on chr20) are used as untyped variants that are imputed by local HMM and by BEAGLE. We classified the variants with respect to MAF by separating variants into 4 different MAF ranges: 1) MAF ∈ [0, 0.005] (Very rare), 2) MAF ∈ [0.005, 0.01] (Rare), 3) MAF ∈ [0.01, 0.05] (Uncommon), 4) MAF ∈ [0.05, 0.5] (Common). Before imputing untyped variants, the genotypes are phased using Eagle2^51^. We use the parameters *(l*_*w*_, *N*_*e*_, *l*_*c*2*t*_, *n*_*tag*_) = (0.5,10^3^, 0.02, 1000) for validation of accuracy. Figure 4 shows the *R*^2^ and non-reference genotype concordance distribution for the variants on chromosome 19 (Fig. 4a,b) and chromosome 20 (Fig. 4c,d). The imputation of variants in the MAF range of common and uncommon are comparable with the baseline imputations of BEAGLE with less than 1% different in accuracy between baseline and local HMM. For rare and very rare variants, BEAGLE outperforms with more than 2 percent accurately in R2. The non-reference genotype concordance is less than 1 percent for these categories. These results indicate that local HMMs can potentially provide utility for uncommon and common variants (i.e., MAF>1%).

**Figure 4.**
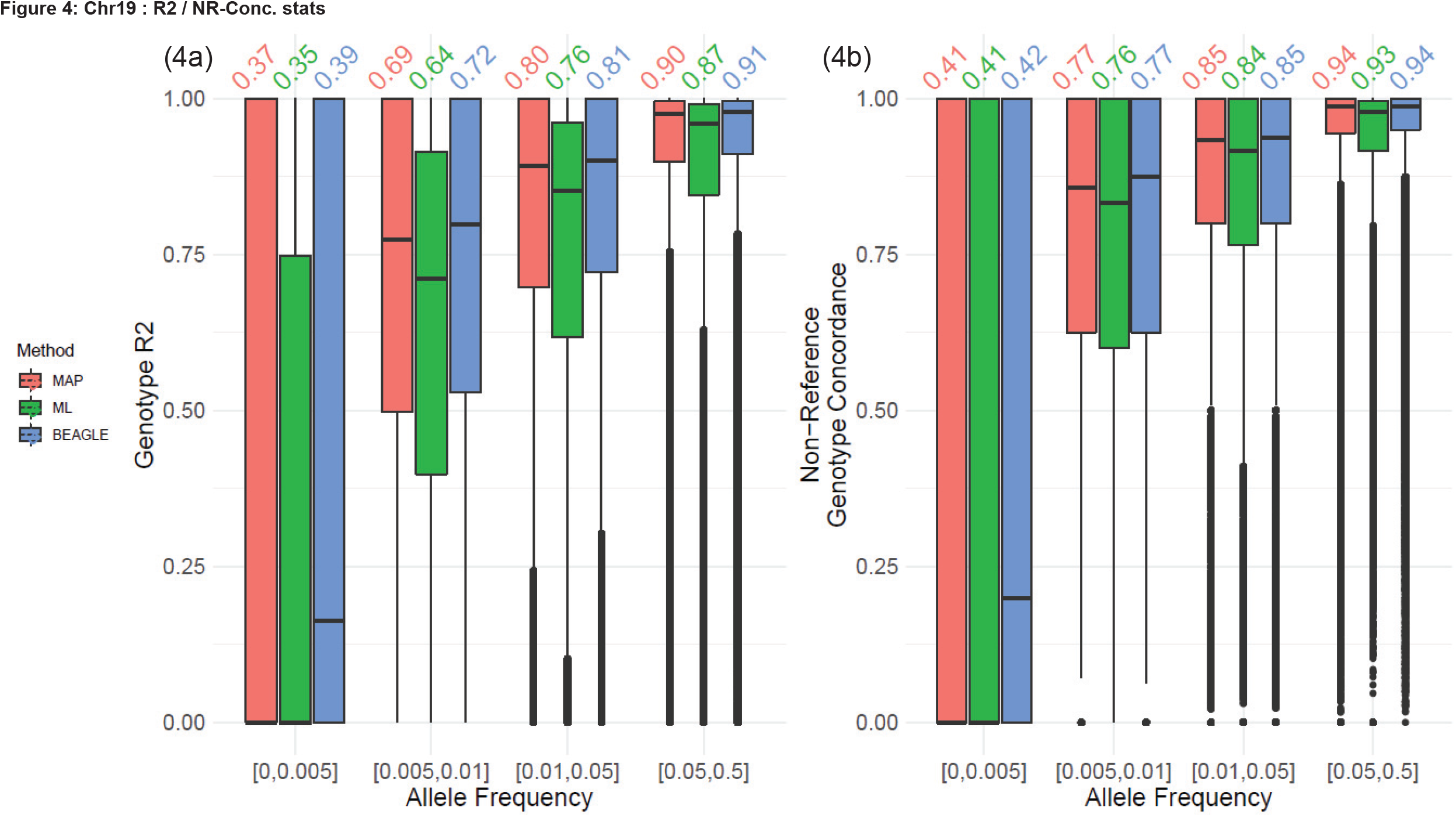
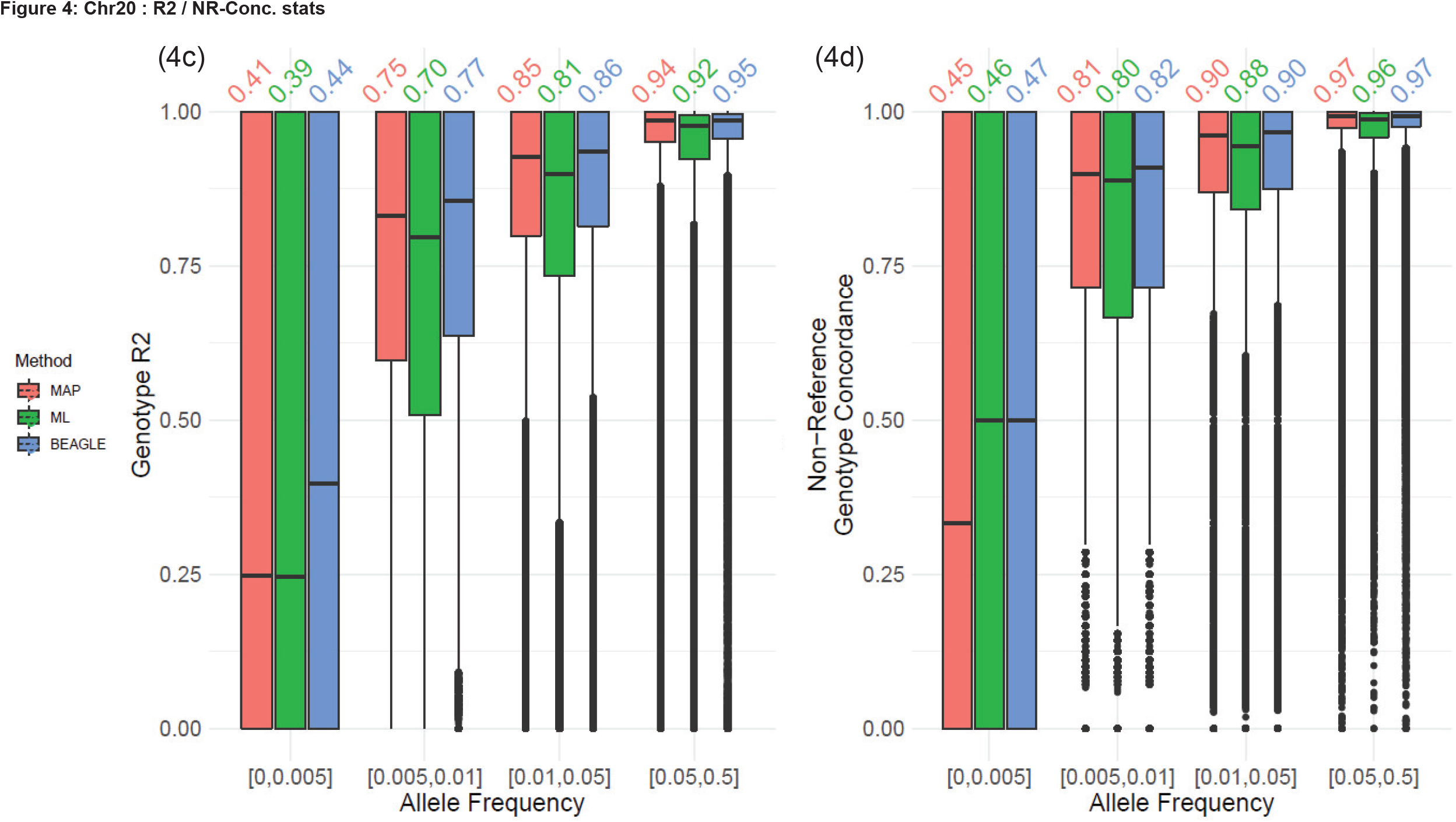
**(a)** Distribution of genotype *R*^2^ and **(b)** non-reference genotype concordance for the untyped variants on chromosome 19. The variants are stratified with respect to minor allele frequency (MAF) as shown on the x-axis. **(a)** Distribution of genotype *R*^2^ and **(b)** non-reference genotype concordance for the untyped variants on chromosome 20.

## Discussion

We analyzed the feasibility of imputing variants using HMMs that are computed on locality of the target variants, i.e. untyped variants. There are several advantages of focusing exclusively to the locality of an untyped target. First, the computations can be parallelized and performed at a much smaller scale without the need of large number of untyped variants. Second, the evaluation of assessment of the local HMM accuracy can provide biological insight into the haplotype structure and imputability estimates^58^. Third, the local models can be run in isolation from other parts of the genome. This way, the imputation algorithms can be re-designed for other tasks. For instance, recently developed privacy-aware imputation methods make extensive use of the locality-based models. Our results provide insight into the design of secure imputation algorithms. Also, our study provides evidence that HMM-based imputation methods can be designed with a locality-based approach.

It should be noted that the default parameters does not provide the optimal performance that can be achieved using local HMMs. For instance, we did not evaluate the impact of increasing *l*_*w*_ while the maximum number of tag variants is kept constant. This would still constitute a local HMM model since the maximum number of surrounding tags is constrained. In other words, this would keep the computational requirements constant but it would enable local HMM to assess large haplotype blocks. In addition, the locality windows can be implemented in different ways. For example, the tags can be filtered with respect to the smallest genetic distance, i.e., we can remove the tag variants that are close to each other.

The main limitation of the local-HMM methods that are evaluated here is the lower accuracy for rare variants, especially for the variants with MAF lower than 1%. Our results show that the performance can be improved by extending the local windows to include more variants. This is reasonable since longer windows enable the resolution of the rare haplotypes more accurately than shorter windows. From a utility perspective, we observed that most of the downstream analyses, such as genomewide association studies (GWAS) impose thresholds on the well above 1%. For instance, even high powered GWAS studies impose thresholds at 2-5% on the MAF of the variants to provide enough power for detecting phenotype-genotype associations^59^. Also, even the state-of-the-art HMM methods do not provide the imputation accuracy for low MAF variants that is necessary for the downstream analyses. In addition, these rare variants tend to be population-specific^60^ and usage of population specific panels can enable more accurate performance. Thus, the local HMMs can be used to impute variants for downstream tasks with MAF values that are utilizable for studies such as GWAS.

## Methods

We present the computational details of the ML and MAP estimation from the local-HMM.

### Description of the Local-HMMs

LoHaMMer computes MAP and ML estimates on the typed tag variants, i.e., keeps track of haplotype paths that are passing through only the typed variants. We assume that the genotypes are phased and the genotype matrix is denoted by 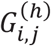, which stands for the allele on parental copy *h* for individual *i* and the variant at index *j*. The parental copy has two values *h* ∈ {0,1}, indicating the paternal and maternal haplotypes (or vice versa). 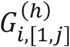 indicates the sequence of alleles for *i*^*th*^ individual for variants between *1* and *j*, i.e., 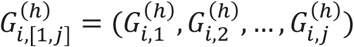. The alleles for each variant can have 2 values, 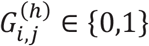, denoting reference allele and alternate alleles. 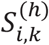 denotes the HMM state at the variant *k* for *i*^*th*^ individual. The states correspond to the indices of haplotypes in the phased reference genotype panel, i.e., 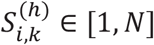. We denote the indices of the untyped variants with *j*_*Ø*_, which is the set of variant indices (i.e., *j* < *V*) for which the genotypes are missing.

#### Variant Subsampling

Given the maximum number of typed variants (or tag variants), 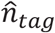, LoHaMMer first identifies all the variants in the current window, which is of length *l*_*w*_. Given that 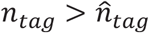 is the total number of variants, LoHaMMer takes every 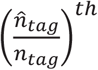 variant to select 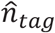 in the window. If *n*_*tag*_ is smaller than 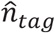, all the tag variants are used for imputation. To simplify the presentation, we assume that the variant indexing is based on the subsampled variant list.

#### Posterior Probability Estimation by Forward-Backward Algorithm (MAP Method)

The MAP algorithm relies on computation of forward and backward matrices. Given individual *i* and haplotype *h*, the forward probability is formulated as:

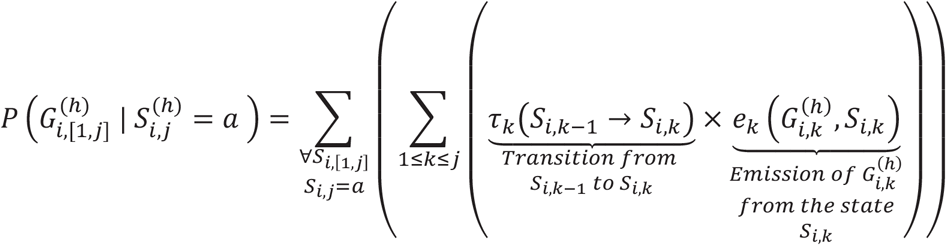

where 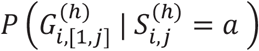 denotes the forward variable, which is the total probability of all state sequences and the emissions from the state sequences *S*_*i*,[1,*j*]_ that emit the allele sequence 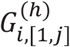 with the constraint that the last state at variant *j* is *a*, i.e., 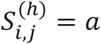. The forward variable matrix can be computed for all variant positions and all states using recursion:

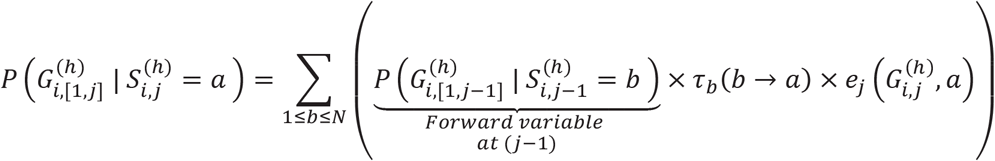

where the forward variable at variant *j* is computed using the forward variable at position *(j* − 1). The boundary condition is defined at the first nucleotide:

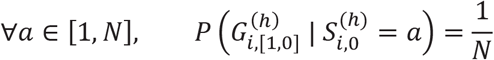

which indicates that the state at the first variant is uniformly distributed among all states, i.e. there is no preference between haplotypes that initiate the HMM. This boundary condition is sometimes described by introducing a special state named the “start state”.

The backward probability is formulated as:

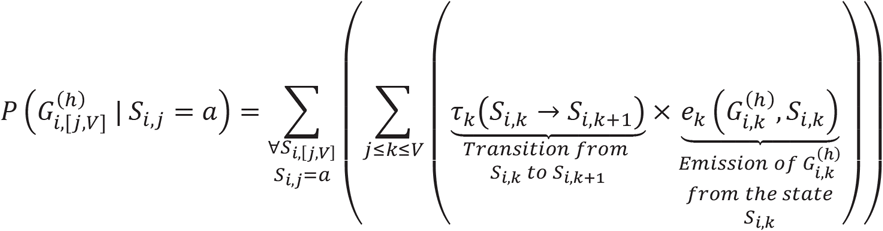

denotes the backward matrix for *i*^*th*^ individual’s haplotype *h*, and the total probability over all the state sequences, *S*_*i*,[*j,V*]_, that emit the allele subsequence 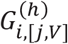 with the constraint that the first state at variant *j* is *a*, i.e., 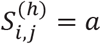. The backward variable can be computed using a simple recursion relationship, similar to the forward variable:

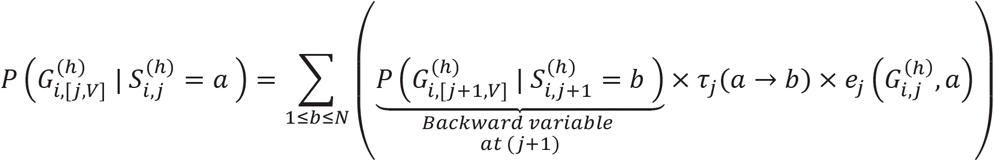

with the boundary conditions:

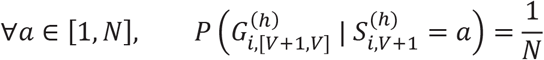

which indicates that the haplotypes are uniformly distributed at the end of the allele sequence.

#### Computation of the Posterior Probability for Untyped Variants

The forward and backward variables are used for inferring the probability of the observing alleles 0 and 1 at the untyped variants. To estimate the allele probabilities of an untyped variant at index *j* ∈ *j*_Ø_, LoHaMMer identifies the two consecutive typed variants that are closest to the variant *j*. Using the nearest tags, LoHaMMer uses an approach similar to BEAGLE to estimate the path that passes along the untyped variant:

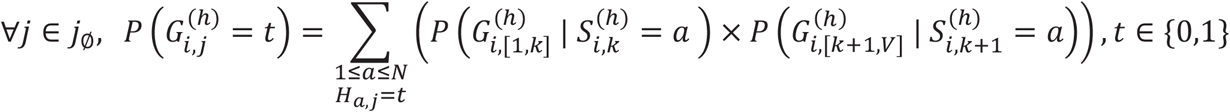

where *j*_Ø_ indicates the untyped variant indices in the genotype matrix, and *k* is the variant index such that variants at *k* and *k* + 1 are the closest typed tags to untyped variant *j*. The allelic probabilities from the parental copies are normalized and combined to generate a final genotype probability for the 3 possible genotypes:

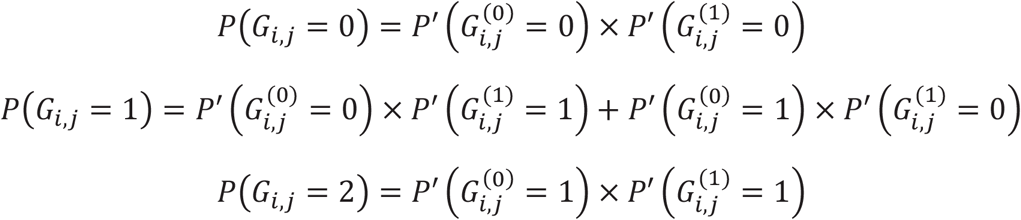

where 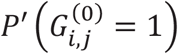 denotes the normalized probabilities that are in the range [0,1]. These are computed as following

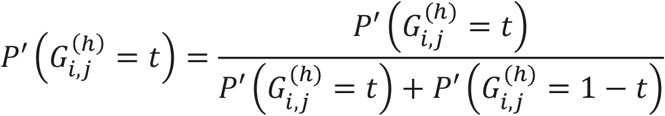

where the normalization is performed over the two possible allelic probabilities for the parental copy *t*.

#### Maximum-Likelihood Haplotype Path Estimation by Viterbi Algorithm (ML Method)

Similar to the forward matrices, the ML method keeps track of the maximum scoring matrix at each typed tag for every possible haplotype state:

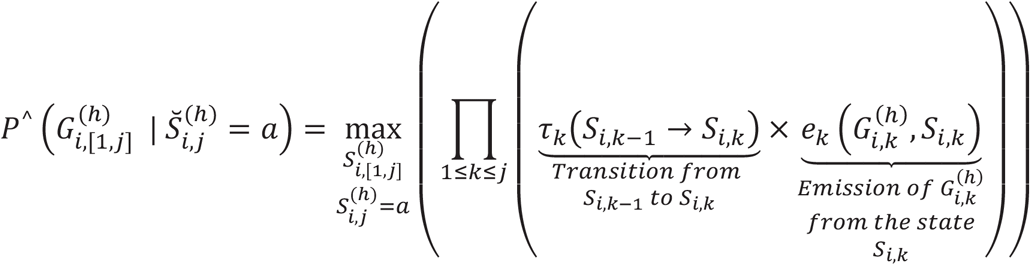

where 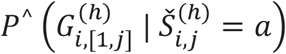 indicates the probability of the typed allele sequence 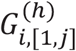 from the most likely state sequence 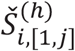 with additional constraint of 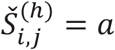. This path is the ML path that LoHaMMer uses to infer the most likely haplotype mosaic that emits the typed allele sequence. This array is computed using the following recursion relationship:

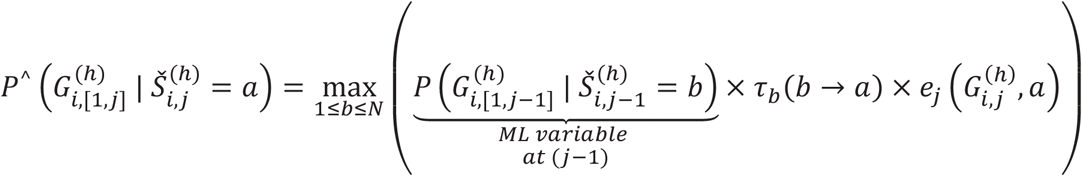

LoHaMMer computes the ML matrix using this recursion relationship for every typed tag variant from left to right for all the haplotypes with the boundary condition:

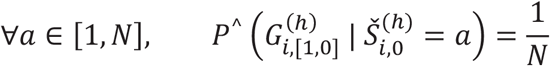

As for the forward and backward matrices, the ML matrix is computed over the typed tag variants.

After computing the ML matrix, LoHaMMer traces back the ML matrix to identify the optimal state sequence i.e., the optimal set of haplotypes that emits the full allelic sequence:

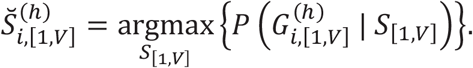

After the optimal state is assigned, LoHaMMer assigns the alleles to the untyped variants similar to the MAP method. For the untyped variant at index *j*, LoHaMMer identifies the closest typed tag variant and assigns the allele based on the state on the tag variant:

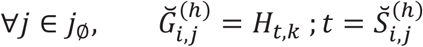

where *k* is the tag variant that is closest to the untyped variant at *j*.

#### Haplotype Clustering in Blocks of Variants

The recursion relationships for ML and forward-backward variants indicate that it is necessary to perform a summation (or a maximum operation) over all the haplotypes in the reference panel, for every typed tag variant. This computation can become quickly intractable. Similar to the previous methods, LoHaMMer clusters the haplotypes, computes each MAP and ML arrays over the clusters of reference haplotypes to minimize the number of redundant operations. The clustering increases the efficiency substantially because (1) the number of unique haplotypes over short stretches increase much slower compared to the number of haplotypes, (2) the transition probabilities between states depends only on the self-transition and recombinations. These optimizations are extensively described in previous methods.

We will briefly describe the usage of clustering for computation of ML arrays. LoHaMMer selects a number of variants that will be used to cluster the reference haplotypes, by default the block length is selected to be 10 variants. Given a local window, LoHaMMer divides the window into blocks of 10 variants. Next the reference haplotypes on each block are clustered such that each cluster corresponds to a unique sequence of 10 alleles, corresponding to 10 variants in the block. Next, for each cluster, the ML variable is computed as the maximum of the ML variable over the haplotypes in the cluster. Since the clusters share the allelic sequence exactly, ML variables for the clusters are computed at the cluster-level using the recursion relatonships over the 10 variants in the block. After cluster-level ML variables are computed for each cluster, LoHaMMer assigns the ML variable to each haplotype from their corresponding cluster-level ML variables.

#### Numerical Stability

The transition and emission probabilities are smaller than 1 and they are multiplied with each other over all transitions and emissions. Thus, the ML variable and MAP variables may overrun or underrun the numerical precision. To get around these numerical stability isssues, LoHaMMer can perform the computations in the logarithmic domain or it scales the ML and forward-backward variables by a scaling factor. For the logarithmic domain computations, LoHaMMer keeps every value as logarithms. In logarithmic domain, a multiplication is converted to a summation and this is convenient since the overflow is virtually impossible. However, we observed that the approximate summation in logarithmic domain requires numerous slow operations and causes big decreases in speed. Therefore, LoHaMMer uses a linear scaling value by default. For this, LoHaMMer multiplies every array value by a constant scaling factor. We observed by trial-and-error that scaling factor of exp (0.2) enables minimal number of underflow or overflow issues. LoHaMMer keeps track of any overflow and underflow at each computation step. If an array value becomes too high or too low, the values are re-scaled to ensure numerical stability.

### Computation of Accuracy Metrics

#### Genotype *R*^2^

The genotype *R*^2^ is computed as the squared value of the Pearson correlation coefficient between the known and imputed genotype for each variants. For the *j*^*th*^untyped variant the genotype *R*^2^ is computed as:

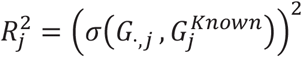

where *σ* indicates the Pearson correlation of the genotype value.

#### Non-Reference Genotype Concordance

The concordance is simply the overlap between the genotypes that are known to be non-reference:

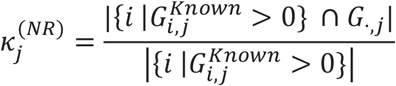

where 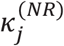 denotes the non-reference concordance between the known non-reference genotypes of variant *j* and the imputed variants over all individuals.

### Data Sources

The 1000 genomes project genotypes are downloaded from NCBI ftp data portal. The Illumina Duo v3 variants are extracted from the array’s documentation available at: ftp://webdata2:webdata2@ussd-ftp.illumina.com/MyIllumina/c1859532-43c5-4df2-a1b4-c9f0374476d2/Human1M-Duov3_H.csv. The variants in The 1000 Genomes Project that overlap with the variants on the array’s tag variants are used as the typed variants.

